# NAD Metabolism Drives CTCF-Cohesin Chromatin Regulation in EBV-Transformed B-Cells

**DOI:** 10.64898/2026.11.01.763250

**Authors:** Davide Maestri, Rachel Sklutuis, Lisa Beatrice Caruso, Sarah Preston-Alp, Martina Gatto, Samantha S. Soldan, Pyounghwa Park, Lauren Melick, Laura Corveleyn, Simone Sidoli, Andrew Kossenkov, Aaron R. Goldman, Wujuan Zhang, Hsin-Yao Tang, Giorgia Napoletani, Yifei Liao, Paul M. Lieberman, Benjamin E. Gewurz, Italo Tempera

## Abstract

Chromatin architecture regulates gene expression, and it is maintained by architectural proteins such as CTCF and the cohesin complex. In Epstein–Barr virus infected cells, viral chromatin organization is stabilized through the interaction between PARP1 and CTCF, ensuring maintenance of latent gene expression. EBV⁺ B cells exhibit elevated PARP1 activity, which imposes a substantial metabolic burden due to NAD⁺ consumption by PARP1 itself. We demonstrate that efficient EBV infection and B-cell transformation rely on the NAD salvage pathway to sustain PARP1 activity and CTCF-cohesin chromatin folding. Pharmacologic inhibition of the rate-limiting enzyme NAMPT with FK866 phenocopies the chromatin architectural disruption induced by PARP1 and changes in NAD levels affects CTCF/cohesin chromatin architectural remodeling and loop extrusion. Moreover, NAMPT inhibition reduces tumor burden in EBV⁺ xenograft models. These findings identify a critical metabolic dependency linking NAD⁺ homeostasis to CTCF/cohesin mediated chromatin architecture and reveal a therapeutic vulnerability in EBV-associated malignancies.

## Introduction

The three-dimensional (3D) organization of chromatin within the nucleus plays a critical role in regulating diverse genomic functions, including gene transcription, DNA replication, and cell differentiation^1–3^. This higher-order genome architecture arises from a complex network of spatial interactions that bring together genomic regions separated by large distances on the linear DNA sequence. Central to these interactions are chromatin loops, which can span from several kilobase pairs (kbp) to megabase pairs (Mbp). Formation and stabilization of chromatin loops largely depend on the cooperative action of the cohesin complex and the architectural protein CTCF^4,5^. The cohesin complex is a ring-shaped, ATP-dependent protein assembly composed of structural maintenance of chromosomes (SMC) subunits SMC1A and SMC3 linked together by the kleisin subunit RAD21, and is further stabilized and regulated by auxiliary proteins such as STAG1/2 and the loading factor NIPBL^6^. CTCF, a zinc finger DNA-binding protein, acts as a crucial architectural component of chromatin, delineating topologically associating domains and serving as a binding anchor for cohesin-mediated loops, thereby orchestrating precise gene regulation and higher-order chromatin structure^7,8^.

The 3D organization of chromatin has emerged as a critical epigenetic regulator in the infectious cycles of DNA viruses. The cellular architectural proteins CTCF and cohesin have been shown to bind viral genomes and orchestrate chromatin looping in diverse DNA viruses, including Human Papillomavirus (HPV)^9,10^, Herpes Simplex Virus (HSV)^11,12^, Kaposi’s Sarcoma-associated Herpesvirus (KSHV)^13–17^, and Epstein–Barr virus (EBV)^18–23^. EBV infects over 90% of the global population, establishing lifelong latency primarily in B-cells, and is etiologically linked to several malignancies, including Burkitt lymphoma, Hodgkin lymphoma, post-transplant lymphoproliferative disease (PTLD), gastric and nasopharyngeal carcinomas^24,25^. A hallmark of EBV latency is the significant reorganization of host-cell transcriptional programs and chromatin architecture, promoting cellular proliferation while simultaneously repressing immunogenic viral lytic gene expression^18,26–35^. Our recent work indicates that this extensive epigenetic remodeling is mediated, in part, by poly-(ADP-ribose)-polymerase-1 (PARP1), an NAD⁺-dependent chromatin-modifying enzyme essential for EBV-driven gene expression and B-cell immortalization^19^. PARP1 utilizes NAD⁺ as a substrate to catalyze the poly(ADP-ribosyl)ation of target proteins, thereby altering their functions. Specifically, PARP1 modifies CTCF through poly(ADP-ribosyl)ation, influencing its chromatin-binding affinity and insulation functions^36^. In EBV-infected B-cells, PARP1 activity is required for effective CTCF recruitment and subsequent chromatin looping across the EBV genome. Pharmacological inhibition of PARP1 destabilizes CTCF binding and viral chromatin organization, leading to heterochromatinization of the EBV genome and repression of viral latent gene expression^18–20,37,38^.

Continuous PARP1 activity in EBV-infected cells imposes a substantial metabolic burden due to the consumption of NAD⁺ during each poly(ADP-ribosyl)ation event. How EBV-transformed B-cells sustain this heightened demand for NAD⁺ remains incompletely understood. Cellular NAD⁺ homeostasis is maintained through three primary biosynthetic pathways: the de novo kynurenine pathway from tryptophan, the Preiss–Handler pathway from nicotinic acid, and the salvage pathway, which recycles nicotinamide via the rate-limiting enzyme nicotinamide phosphoribosyltransferase (NAMPT)^39^. Recent evidence indicates that during the initial stages of EBV infection, indoleamine-2,3-dioxygenase 1 (IDO1)-dependent activation of the de novo pathway supports a transient surge in NAD⁺ synthesis^40^; however, whether this mechanism alone is sufficient to meet the persistent NAD⁺ demands of latently infected, proliferating B-cells is unclear. In addition, given that NAD levels may affect the PARP1–CTCF axis, the extent to which EBV manipulates NAD⁺ metabolism as part of its epigenetic control of host and viral chromatin, especially via CTCF-cohesin–mediated looping, remains unknown.

Here we show that upon infection of B-cells, EBV co-opts the NAD salvage pathway to ensure a continuous supply of NAD⁺ that fuels PARP1 activity, shapes host and viral chromatin topology, and sustains oncogenic transcriptional programs. Metabolomics revealed a marked accumulation of NAD⁺ and nicotinamide — and a reciprocal decrease in quinolinic acid — in primary B-cells after infection, indicating preferential engagement of the NAD salvage pathway. Proteomic profiling corroborated these findings: NAMPT was robustly upregulated while IDO1 was suppressed. Upon infection, pharmacological inhibition of NAD salvage pathway activity with FK866 abrogated both NAD⁺ induction and the outgrowth of EBV+ lymphoblastoid cells. In addition, here we report that the functional integrity of CTCF and cohesin-mediated chromatin looping critically depends on the availability of NAD⁺. Importantly, depletion of NAD⁺ levels affects CTCF and cohesin occupancy across the genome and interferes with proper chromatin loop extrusion, resulting in genomic reorganization, altered transcriptional programs, and compromised viral gene expression. These findings reveal NAD⁺ as an essential metabolic determinant of chromatin architecture, directly linking cellular metabolism to 3D genome organization. Beyond viral infection, our study provides important insights into an underappreciated metabolic regulation of CTCF-cohesin interactions, with broad implications for genome stability, cellular differentiation, and cancer.

## Materials and methods

### Cell culture and treatment

Cell lines were maintained in a humidified incubator containing 5% CO_2_ at 37°C. LCL cells cell lines were cultured in RPMI 1640 supplemented with Fetal Bovine Serum at a concentration of 15%, supplemented with 1% penicillin-streptomycin and plasmocin at a concentration 1:500 (Invivogen ant-mpp). Mutu1 cell lines were cultured under the same conditions except for the concentration of FBS supplemented to the media (10%).

B-cells were negatively selected from PBMCs using EasySep^TM^ Human B Cell Isolation Kit (StemCell Technologies 100-0971) as per manufacturer’s protocol. Cells were then counted and used for experiments carried out before infection. The remaining cells were spun down 600g 3 minutes and resuspended in RPMI 1640 supplemented with HyClone^TM^ Fetal Bovine Serum at a concentration of 15% (Cytiva SH30071.03) and infected with EBV (B95.8 strain).

PARP1 inhibitor (BMN673, Selleck Chemicals Catalog No.S7048) was used at a concentration of 200 nM and cells were treated for 72 hours.

NAMPT inhibitor (FK866, Selleck Chemicals Catalog No.S2799) was used at a concentration of 2.5 nM for 72-hour treatment.

NAMPT PROTAC (NAMPT degrader-2, MedChemExpress catalog No. HY-152154) was used at a concentration of 50 nM for 72-hour treatment with retreatment every 24 hours.

NMN (Fosribnicotinamide, MedChemExpress catalog No. HY-F0004) was used at a concentration of 100 uM for 72-hour treatment with retreatment every 24 hours.

### Western Blot

Protein concentration measurement and western blotting were carried out as previously described^18^ using a bicinchoninic acid (BCA) protein assay (Pierce, Cat. No. 23227). Membranes were incubated overnight at 4°C with primary antibodies against LMP1 (Abcam, Cat. No. ab78113), MYC (Proteintech Cat. No. 10828-1-AP), EBNA2 (Millipore, Cat. No. MABE8), NAMPT (Thermo Fisher Scientific, Cat. No. MA544309), PARP1 (Invitrogen, Cat. No. MA3-950), PAR (RD, Cat. No. 4335MC100) and β-Actin (Cell Signaling Technology 12262S) as per manufacturer’s recommendation.

Quantification was performed using iBright Analysis Software version 5.3.0 (Thermo Fisher Scientific).

### Flow Cytometry and Cell Sorting

Cells were washed with DPBS and stained with Zombie NIR Fixable Viability Dye (BioLegend, Cat. No. 423105) at a 1:1000 dilution in DPBS for 20 minutes at room temperature, protected from light. Following viability staining, cells were washed and incubated with fluorophore-conjugated antibodies against CD19 (APC; BD Pharmingen, Cat. No. 555415) and CD23 (PE; BD Pharmingen, Cat. No. 555711) at 1 μL per 1 × 10⁵ cells for 30 minutes at room temperature in the dark. Cells were subsequently washed and resuspended in sorting buffer for analysis.

Flow cytometric analysis was performed on live (Zombie NIR-negative) cells. For cell sorting experiments, B cells were gated as CD19⁺ and separated into CD23hi and CD23⁻ populations. Sorted populations were collected for downstream analyses.

Cell proliferation was assessed using CellTrace Violet (Thermo Fisher Scientific, Cat No. C34557) staining and flow cytometry. Cells were labeled with CellTrace Violet according to the manufacturer’s instructions, washed to remove excess dye, and cultured under the indicated experimental conditions. Following treatment, cells were harvested and analyzed by flow cytometry to assess dye dilution as a measure of cell division. Proliferation was quantified by comparing CellTrace Violet fluorescence intensity across experimental conditions.

Cell-cycle distribution was assessed by propidium iodide (PI) staining and flow cytometry. Following treatment, cells were harvested, washed with phosphate-buffered saline (PBS), and fixed in cold 70% ethanol. Fixed cells were washed and incubated with PI staining solution containing RNase A to stain DNA and eliminate RNA-associated signal. Samples were analyzed by flow cytometry, and DNA content was used to determine the proportion of cells in the G0/G1, S, and G2/M phases of the cell cycle.

### NAD Quantification

Intracellular NAD levels were quantified using the NAD/NADH-Glo™ Assay (Promega Cat. No. G9071) according to the manufacturer’s instructions with minor modifications. Cells were harvested at the indicated time points, and cell numbers were determined either by trypan blue exclusion or during flow cytometric cell sorting, as indicated for each experiment. Cell lysates were prepared from collected cells using the reagents supplied in the kit and transferred to white-walled 96-well plates for analysis.

To selectively measure NAD, endogenous NADH was degraded prior to detection using the NAD-specific sample preparation procedure recommended by the manufacturer. Briefly, lysates were incubated under alkaline conditions to decompose NADH while preserving NAD, followed by neutralization. The NAD/NADH-Glo™ Detection Reagent was then added, and luminescence was measured using a plate reader according to the manufacturer’s protocol.

Luminescence values were normalized to the corresponding cell number for each sample to account for differences in cell abundance. Relative NAD levels were subsequently normalized to the PBS-treated control group and are presented as relative NAD abundance.

### Dot Blot

Dot Blot was carried out as per Abcam protocol. Briefly, three biological replicates of 2 × 10^6^ LCL cells per treatment group (with or without 2.5 nM FK866 for 72 h) were extracted using radioimmunoprecipitation assay (RIPA) lysis buffer (50 mM Tris–HCl, pH 7.4, 150 mM NaCl, 0.25% deoxycholic acid, 1% NP-40, 1 mM EDTA; Millipore, Cat. No. 20–188) supplemented with 1X protease inhibitor cocktail (Sigma-Aldrich, Cat. No. P8340-5ML) and 1× PARG inhibitor (PDD00017273, Selleck Chemicals, Cat. No. S8862). Protein concentration was measured using a bicinchoninic acid (BCA) protein assay (Pierce, Cat. No. 23227) and 10 μg of protein were diluted in 50 μl of water and loaded onto the activated nitrocellulose membrane (Bio-Rad Cat. No. 1620113). Red Ponceau staining (Ponceau S Staining Solution, Cell Signaling, Cat. No. 59803) was used as a loading control. Membrane was blocked in 5% milk TBS-T and then incubated with anti-PAR antibody (R&D Systems, Cat. No. 4335-MC-100) as per manufacturer’s recommendation. Membrane was washed 5 mins three times, incubated for 1 h with rabbit anti-mouse IgG-HRP (Jackson ImmunoResearch), at a dilution of 1:10000 and detected by enhanced chemiluminescence (SuperSignal West Dura Extended Duration Substrate, Cat. No. 34075).

### Digital Droplet PCR (ddPCR)

DNA was extracted from FK866 and vehicle treated cells using the Qiagen DNeasy blood and tissue kit (Qiagen, Hilden, Germany) according to the manufacturer’s instructions. DNA concentrations were measured using the NanoDrop 2000 Spectrophotometer (ThermoFisher, Waltham, MA) prior to equalization of each sample to 30 ng/µl DNA, followed by HIND III digestion (New England Biolabs, Ipswitch, MA). DdPCR (Bio Rad QX200, Ipswitch, MA) was performed using the following primer probe sets to calculate EBV copy number per cell: EBV-LMP1 primers and FAM/MGB probe (Forward: AAGGTCAAAGAACAAGGCCAAG, Reverse: GCATCGGAGTCGGTGG ; Probe:FAM-AGCGTGTCCCCGTGGAGG; and Ribonuclease P protein subunit 30 (RPP30) primers and VIC/MGB probe (Forward: GATTTGGACCTGCGAGCG, Reverse: GCGGCTGTCTCCACAAGT; Probe: VIC-CTGACCTGAAGGCTCT)^41,42^.

### RNA extraction and RNA-seq

Total RNA from LCL cell lines per treatment group was isolated from 2×10^6^ cells using a Direct-zol RNA Kit (Zymo Research, cat. No. R2050) according to the manufacturer’s protocol. RNA samples were submitted to the Wistar Institute genomics core facility for RNA quality control and sequencing library preparation using the SENSE mRNA-Seq Library Prep Kit V2 (Lexogen) to generate Illumina-compatible sequencing libraries according to the manufacturer’s instructions. Paired-end reads of 150 bp were obtained using an Illumina HiSeq 2500 sequencer.

### RNA-seq Analysis

RNA-seq data was aligned using STAR^43^ against hg19 version of the Human genome and all unaligned reads were then aligned against NC_007605.1 version of EBV genome and *featureCounts* software^44^ was used to estimate raw read counts for Human and EBV genes. DESeq2^45^ was used to perform differential analysis. DeepTools^46^ was used for data visualization. To compare different conditions, alignment files were normalized using the reciprocal of the size factors calculated in DESeq2^45^ and inputted as the *--scaleFactor* argument of the *bamCoverage* function of deepTools^46^. For visualization purposes all replicates for the same condition were merged. Ingenuity Pathway Analysis (IPA)^47^ was used to perform gene ontology and to identify the upstream regulators of differentially expressed genes. Heatmaps for differentially expressed genes were generated using *pheatmap* R package^48^. Datasets are available in Gene Expression Omnibus see Data Availability section for accession number.

### dCas9–KRAB-mediated CRISPR interference

To silence NAMPT enhancer we used CRISPRi. Briefly, sgRNAs (sg1: CAGCCTCAAACAGTGTCATATGG, sg2: GCCAAACATTAATTATATCTTGG, sg3: CTAAGTTGTTTGACATAGCAGGG and sgCTRL: ATTTCGCAGATCATCGACAT) were cloned into the lentiGuide-Puro plasmid (Addgene #52963). Then GM12878 cell lines stably overexpressing the dCas9–KRAB-MeCP2 fusion protein were transduced with lentiviruses containing the sgRNAs and then selected with 3mg/mL of puromycin for 5 days.

### Chromatin immunoprecipitation assays

#### Chromatin immunoprecipitation sequencing (ChIP-seq)

Chromatin immunoprecipitation with next-generation sequencing (ChIP-seq) was performed as previously described^19^. Briefly, 50×10^6^ cells per immunoprecipitation were collected and fixed with 1% formaldehyde for 15 min and then quenched with 0.25 M glycine for 5 min on ice. After 3 washes with 1X PBS, pellets were resuspended in 10 mL each of a series of two lysis buffers and resuspended in 1mL of the third lysis buffer before fragmentation in Covaris ME220 Ultrasonicator (peak power 75, duty factor 25, cycles/burst 1000, average power 18.8, time 720 sec) to generate chromatin fragments roughly 200–500 bp in size. Chromatin was centrifuged to clear debris and a 1:20 of this cleared chromatin was kept as standard input for comparison against immunoprecipitations. Chromatin was incubated rotating at 4°C 1h with 20 µg CTCF (Active Motif 61311), 25 µg SMC3 (Bethyl A300-060A) and 25ug NIPBL (Bethyl A301-779A), then chromatin– antibody complexes were precipitated using 250 µL of Dynabeads Protein A (ThermoFisher, product No. 10001D) incubated rotating at 4°C overnight. DNA was purified using Zymo ChIP DNA Clean & Concentrator Kit (Zymo Research, product No. D5205). Libraries for sequencing were made using NEBNext Ultra II DNA Library Prep Kit (New England Biolabs, product No. E7103) and sequenced on the Illumina HiSeq 2500.

### ChIP-seq analysis

Reads were mapped against the hg19 version of the human genome and the human gammaherpesvirus 4 (HHV4) NC_007605.1 genome assembly using Burrows-Wheeler Aligner (BWA)^49^. We used MACS2^50,51^ software package to call peaks using input samples as control. deepTools^46^ was used for data visualization.

For comparisons, alignment files were read-depth normalized and replicates were merged for visualization purposes.

For transcription factor binding motif analysis, HOMER^52^ *findMotifsGenome.pl* function was used using default options. HOMER *annotatePeaks.pl* function was used to annotate peaks. DESeq2^45^ was used to do differential analysis of ChIP-seq peaks. Average plots were obtained using seqMINER^53^ and plotted using *ggplot2* R package^54^.

ChIP-seq data were deposited for public access at Gene Expression Omnibus see Data Availability section for accession number.

### CUT&Tag

CUT&Tag was performed as per EpiCypher protocol. Briefly, 100,000 cells per condition were crosslinked using 0.1% formaldehyde for 1 minute, then crosslinking was quenched using 0.125M glycine. Cells were spun down 600g 3 minutes, and nuclei were extracted by resuspending cells in Nuclei Extraction Buffer (20 mM HEPES-KOH pH 7.9, 10 mM KCl, 0.1% Triton X-100, 20% Glycerol, 1X Protease Inhibitor without EDTA, 0.5 mM Spermidine) on ice for 10 minutes. Nuclei were spun 600g for 3 minutes and resuspended in Nuclei Extraction Buffer and added to 10uL of Activated ConA beads (Bead Activation Buffer: 20 mM HEPES pH 7.9, 10 mM KCl, 1 mM CaCl_2_, 1 mM MnCl_2_) and incubated on nutator for 10 minutes. Beads were then collected on a magnet and 50uL/reaction of fresh ice-cold Antibody Buffer 150 (20 mM HEPES-KOH pH 7.9, 150 mM NaCl, 1% Triton X-100, 0.05% SDS, 1X Protease Inhibitor without EDTA, 0.5 mM Spermidine, 0.01% Digitonin, 2 mM EDTA) was added to the beads. 1ug of H3K27ac antibody (Active Motif 39133), or 1 ug of IgG antibody (CST 66362S) was added to the reaction and incubated o/n on a nutator at 4°C with cap elevated.

Next day a Secondary Antibody Master Mix was prepared by adding 50uL of Digitonin 150 Buffer (20 mM HEPES-KOH pH 7.9, 150 mM NaCl, 1% Triton X-100, 0.05% SDS, 1X Protease Inhibitor without EDTA, 0.5 mM Spermidine, 0.01% Digitonin) and 0.5 ug of Rabbit Secondary Antibody (Epicypher 13-1047) per reaction. Beads were collected on a magnet, supernatant was removed and 50uL of the master mix were added to each tube. Tubes were vortexed to mix and incubated for 30 minutes at RT on a nutator. Beads were than washed twice with 200uL of ice-cold Digitonin Buffer 150, resuspended in 50 uL of Digitonin300 Buffer (20 mM HEPES-KOH pH 7.9, 300 mM NaCl, 1% Triton X-100, 0.05% SDS, 1X Protease Inhibitor without EDTA, 0.5 mM Spermidine, 0.01% Digitonin) and 2.5 uL of pAG-Tn5 (Epicypher 15-1117) was added to each reaction. Samples were incubated for 1 hour at RT on nutator with caps elevated. Beads were washed twice with ice-cold Digitonin300 Buffer and resuspended in 50 uL of cold Tagmentation Buffer (Digitonin300 Buffer, 10mM MgCl_2_). Reactions were incubated for 1 hour at 37C in a thermocycler. Beads were then collected using the magnetic rack, supernatant was aspirated by pipetting, and beads were resuspended in TAPS Buffer (10mM TAPS, pH 8.5, 0.2mM EDTA) by pipetting. After removing TAPS Buffer beads were resusupended in SDS Release Buffer (10mM TAPS, pH 8.5, 0.1 SDS) by vortexing at max speed for 7 seconds. Reactions were then incubated at 58C for 1 hour in a thermocycler to release tagmented DNA. After incubation 15uL of SDS Quench Buffer (0.67% Triton-X 100) were added to each reaction. 2uL of i5 and i7 barcoded primers were added to the DNA together with 25uL of CUTANA High Fidelity 2x PCR Master Mix (Epicypher 15-1018) and PCR was performed as per protocol. DNA was then purified using 1.3X AMPure XP Beads (Beckman-Coulter A63881), eluted in 15uL of 0.1X TE Buffer and quantified using Qubit 1X dsDNA HS quantification kit (ThermoFisher Q33230). Libraries were sequenced on the Illumina HiSeq 2500.

### CUT&Tag Analysis

Reads were mapped against the hg19 version of the human genome and the human gammaherpesvirus 4 (HHV4) NC_007605.1 genome assembly using Burrows-Wheeler Aligner (BWA)^49^. We used HOMER to call peaks using *-type histone* options. HOMER^52^ *annotatePeaks.pl* function was used to annotate peaks. deepTools^46^ was used for data visualization. For comparisons, alignment files for H3K27ac experiments in control and FK866-treated LCLs were read-depth normalized and replicates were merged for visualization purposes. CUT&Tag data were deposited for public access at Gene Expression Omnibus. See Data Availability section for accession number.

### Assay for Transposase-Accessible Chromatin using sequencing (ATAC-seq)

ATAC-seq was performed as described by Grandi et al.^55^. Briefly, 100,000 cells were spun down at 500g for 5 min at 4°C. Cell pellet was resuspended in 50 µL of fresh ice-cold ATAC-seq Lysis Buffer by pipetting up and down three times and incubated on ice for 3 min. Then, 1 mL of fresh ice-cold ATAC-seq Wash Buffer was added to dilute the lysis reagents and tubes inverted five times to mix. Nuclei were spun down at 500g for 10 min at 4 °C. Nuclei pellets were then resuspended in 50 µL of freshly made Transposition Mix by pipetting up and down six times and reactions were incubated at 37°C for 30 min in a thermomixer with 1,000 rpm mixing. The transposition reaction was terminated by adding 250 µL (five volumes) of DNA Binding Buffer from the DNA Clean and Concentrator-5 Kit and mixed well by inversion. Reactions were kept o/n at −20°C. The next day DNA was purified using the Zymo DNA Clean and Concentrator-5 Kit as per protocol. Each sample in the study was assigned to a unique combination of the Adapter 1 (Ad1) and Adapter 2 (Ad2) sequences. Each cleaned-up transposed DNA sample was transferred to a 200 µL PCR tube and mixed with 25 µL of NEBNext Ultra II Q5 2X Master Mix, 2.5 µL of the corresponding Ad1 and 2.5 µL of the corresponding Ad2.

### ATAC-seq Analysis

Reads were mapped against the hg19 version of the human genome and the human gammaherpesvirus 4 (HHV4) NC_007605.1 genome assembly using Burrows-Wheeler Aligner^49^. MACS2^50^ software package was used to call peaks. deepTools^46^ was used for data visualization. For comparisons, alignment files were read-depth normalized and replicates were merged for visualization purposes using samtools^56^. DESeq2^45^ was used for differential analysis of ATAC-seq peaks and HOMER software was used to identify enriched transcription factor motifs in differentially accessible regions. *diffTF* software^57^ was used to assess differential TF binding sites accessibility and function by integrating ATAC-seq data and RNA-seq raw counts.

### Chromatin Conformation Capture Assays HiC assay

Hi-C assay was performed as previously described^19^. Briefly, 5×10^6^ cells per condition were collected for *in-situ* Hi-C. Libraries of total ligation products were produced using Ultralow Library Systems V2 (Tecan Genomics, part no. 0344NB-32) as per manufacturer’s protocol. Libraries were sequenced using the Illumina HiSeq 2500 sequencing platform with paired-end 75bp read length. HiC data was preprocessed using HiC-Pro v2.10.0 pipeline^58^ with default settings using hg19 human genome. HiC matrices for the same condition were combined to get 600M reads per condition and HiCcompare^59^ was used to estimate significance of differential contact based on raw count matrix files at 150kb resolution. *GENOVA* R package^60^ was used to estimate the Relative Contact Probability (RCP), insulation score at CTCF peaks, to generate differential Aggregate Peak Analysis (APA), to perform Aggregate TAD Analysis (ATA), to perform TAD+N analysis and to generate saddle plots for A/B compartments. *cooltools* software package^61^ was used to generate standard APA plots for all chromatin conformation assays. *plotgardener* R package^62^ was used to generate HiC matrix plots. HOMER^52^ was used to detect significant A/B compartment switching and *ggplot2* R package^54^ was used to generate a scatterplot for A/B compartments. To detect stripes from HiC Kight-Ruitz normalized matrices at a resolution of 10kb *stripenn*^63^ software was used. *stripenn compute* with the option *--maxpixel 0.93,0.94,0.95,0.96,0.97,0.98,0.99* was used to detect stripes, while *stripenn score* was used to compute stripiness for every detected stripe in both conditions. The function *stripenn seeimage* was used to plot images of the stripes. The detailed protocol with all minor alterations will be happily supplied by the corresponding author per request. Datasets are available in GEO see Data Availability section for the accession number.

### HiChIP assay

HiChIP assay was performed as per protocol^64^ with minor changes as previously described^18,35^.

Briefly, 10×10^6^ cells per condition were fixed with 1% formaldehyde for 10 min and then quenched with 0.125 M glycine for 5 min at RT. Ligation products were immunoprecipitated with 50 μg H3K27ac antibody (Abcam Cat. No. ab177178). Around 5 μL of Streptavidin C-1 beads were used for biotin pull-down. After the last wash with Tween Wash Buffer, bead-bound DNA was used to prepare libraries with NEBNext Ultra II DNA Library Prep Kit for Illumina (New England Biolabs, Cat. No. E7645L) per manufacturer’s protocol.

Libraries were then deep sequenced using the Illumina HiSeq 2500 sequencing platform.

Data were preprocessed using HiC-Pro v2.10.0 pipeline^58^ with default settings using a combined version of hg19 and EBV NC_007605.1 genomes at 5kb resolution and *hichipper* software^65^ was used to perform restriction site bias modeling and interaction identification using the settings COMBINED/ALL for peak calling from HiChIP samples.

Only statistically significant interactions (FDR < 5%) were kept for downstream analysis.

*diffloop* Bioconductor R package^66^ was used to merge anchors between samples, interactions with less than 5 reads in each condition were discarded and differential loops for each condition were obtained using diffloop edgeR wrapper. Matrices from HiChIP replicates were then merged, read-depth normalized using HiCExplorer^67–69^ and balanced using Knight-Ruiz (KR) normalization and plotted using *plotgardener* R package^62^.

H3K27ac HiChIP anchors were overlapped to SMC3 peaks obtained from previous ChIP-seq experiments using *GenomicRanges*^70^ findOverlaps() function.

### TopoLink assay

TopoLink assay was performed as per Dovetail Genomics protocol. 1×10^6^ cells were used for each condition. Two biological replicates for the control condition were used while three biological replicates were used for BMN673 and FK866 treatment. For each condition and replicate two technical replicates were generated to increase library complexity necessary for loop calling purposes. Technical replicates were then merged computationally after sequencing. The subsequent bioinformatic analysis of the sequencing data was carried out as suggested by Dovetail Genomics (https://topolink.readthedocs.io/en/latest/). *cooltools* software package^61^ was used to generate APA plots and *HiCDC+* R package^71^ was used to call loops. Only loops that had an FDR < 0.05 were kept and those that were present in at least two biological replicates were kept for subsequent analyses.

### Histone extraction and digestion

Histone proteins were extracted from the pellet as previously described^72^ to ensure good-quality identification and quantification of single histone marks. Briefly, histones were acid-extracted with chilled 0.2 M sulfuric acid (5:1, sulfuric acid : pellet) and incubated with constant rotation for 4 h at 4°C, followed by precipitation with 33% trichloroacetic acid (TCA) overnight at 4oC. Then, the supernatant was removed and the tubes were rinsed with ice-cold acetone containing 0.1% HCl, centrifuged and rinsed again using 100% ice-cold acetone. After the final centrifugation, the supernatant was discarded and the pellet was dried using a vacuum centrifuge. The pellet was dissolved in 50 mM ammonium bicarbonate, pH 8.0, and histones were subjected to derivatization using 5 µL of propionic anhydride and 14 µL of ammonium hydroxide (all Sigma Aldrich) to balance the pH at 8.0. The mixture was incubated for 15 min and the procedure was repeated. Histones were then digested with 1 µg of sequencing grade trypsin (Promega) diluted in 50mM ammonium bicarbonate (1:20, enzyme:sample) overnight at room temperature. Derivatization reaction was repeated to derivatize peptide N-termini. The samples were dried in a vacuum centrifuge.

### Sample desalting

Prior to mass spectrometry analysis, samples were desalted using a 96-well plate filter (Orochem) packed with 1 mg of Oasis HLB C-18 resin (Waters). Briefly, the samples were resuspended in 100 µl of 0.1% TFA and loaded onto the HLB resin, which was previously equilibrated using 100 µl of the same buffer. After washing with 100 µl of 0.1% TFA, the samples were eluted with a buffer containing 70 µl of 60% acetonitrile and 0.1% TFA and then dried in a vacuum centrifuge.

### LC-MS/MS Acquisition and Analysis

Samples were resuspended in 10 µl of 0.1% TFA and loaded onto a Dionex RSLC Ultimate 300 (Thermo Scientific), coupled online with an Orbitrap Fusion Lumos (Thermo Scientific). Chromatographic separation was performed with a two-column system, consisting of a C-18 trap cartridge (300 µm ID, 5 mm length) and a picofrit analytical column (75 µm ID, 25 cm length) packed in-house with reversed-phase Repro-Sil Pur C18-AQ 3 µm resin. Peptides were separated using a 30 min gradient from 1-30% buffer B (buffer A: 0.1% formic acid, buffer B: 80% acetonitrile + 0.1% formic acid) at a flow rate of 300 nl/min. The mass spectrometer was set to acquire spectra in a data-independent acquisition (DIA) mode. Briefly, the full MS scan was set to 300-1100 m/z in the orbitrap with a resolution of 120,000 (at 200 m/z) and an AGC target of 5×10e5. MS/MS was performed in the orbitrap with sequential isolation windows of 50 m/z with an AGC target of 2×10^5^ and an HCD collision energy of 30. Histone peptides raw files were imported into EpiProfile 2.0 software^73^. From the extracted ion chromatogram, the area under the curve was obtained and used to estimate the abundance of each peptide. To achieve the relative abundance of post-translational modifications (PTMs), the sum of all different modified forms of a histone peptide was considered as 100% and the area of the particular peptide was divided by the total area for that histone peptide in all of its modified forms. The relative ratio of two isobaric forms was estimated by averaging the ratio for each fragment ion with different mass between the two species. The resulting peptide lists generated by EpiProfile were exported to Microsoft Excel and further processed for a detailed analysis. Proteomics data for Figure 1C were generated by LC-MS/MS as described previously^74^.

**Figure 1:**
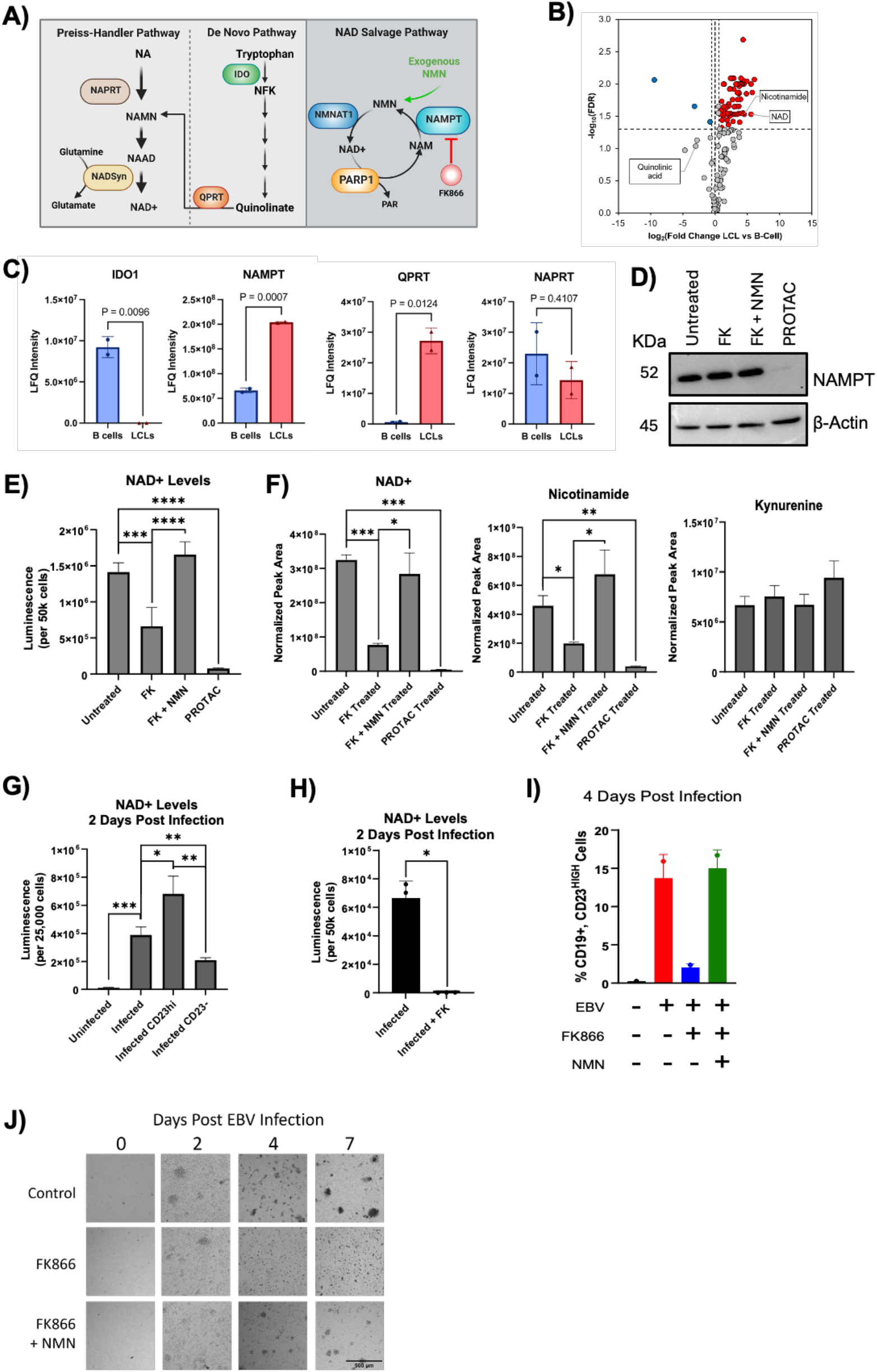
NAD salvage pathway is required for B-cell transformation and is triggered by EBV latent transcription factors. Model of the three metabolic pathways that produce NAD^+^ (**A**). Volcano plot showing differential abundance of metabolites between donor matched LCLs and B-cells (FDR < 0.1) (**B**). Bar plots showing IDO1, NAMPT, QPRT and NAPRT protein abundance between donor matched B-cells and LCLs; p-value determined by student t test on N=3 biological replicates and is indicated on top of each plot (**C**). Western blot showing NAMPT degradation after 50nM NAMPT degrader-2 PROTAC (retreated every day) treatment, but not 2.5nM FK866 treatment **(D**). Bar plot showing NAD+ levels measured by NAD/NADH Glo, a bioluminescent assay to detect NAD+. NADH was first degraded in 50 thousand cells per kit instructions. NAD+ was measured 72 hours after treatment with 2.5nM FK866, 100uM NMN (retreated every day), or 50nM NAMPT degrader-2 PROTAC (retreated every day). Statistical significance was determined using a two-tailed unpaired t-test with Welch’s correction (**E**). Normalized peak area represents the integrated LC-MS peak area following QC Sum normalization to internal standards, providing a measure of relative metabolite abundance across samples. Metabolomic data is from LCLs 72 hours after treatment with 2.5nM FK866, 100uM NMN (retreated every day), or 50nM NAMPT degrader-2 PROTAC (retreated every day). Statistical significance was determined using a two-tailed unpaired t-test with Welch’s correction (**F**). Bar plot illustrating NAD levels (expressed as luminescence over 25×10^6^ cells) between sorted B-cell subpopulations based on CD23 expression levels. Statistical significance was determined using a two-tailed unpaired t-test with Welch’s correction (**G**). Bar plot illustrating NAD levels (expressed as luminescence over 50×10^6^ cells) between Infected and Infected + FK B-cells 2 days post infection. Statistical significance was determined using a two-tailed unpaired t-test with Welch’s correction (**H**). Bar plot showing the percentage of CD19 and CD23 positive cells at 4 days post-infection with B95.8 virus at MOI ∼1 under different treatment conditions (uninfected B-cells, infected B-cells, infected B-cells treated with FK866, infected B-cells treated with FK866 and supplemented with NMN metabolite) (**I**). Microscopic images showing B-cell clumps size at 0, 1, 4 and 7 days post-infection and treated with DMSO, FK866 or FK866 and supplemented with NMN (**I**). Microscopic images showing B-cell clumps size at 0, 1, 4 and 7 days post-infection when treated with 2.5nM FK866, 2.5nM FK866 and supplemented with NMN, or left untreated (**J**).

### Animal studies

All animal experiments were conducted under the Wistar Institute’s approved Institutional Animal Care and Use Committee Protocol #201524 in accordance with the Committee for the Purpose of Control and Supervision of Experiments on Animals guidelines for animal experimentation and the NIH Office of Laboratory Animal Welfare: “PHS Policy on the Humane Care and Use of Research Animals”; the recommendations of the American Association for Accreditation of Laboratory Animal Care (AAALAC): The Guide for the Care and Use of Laboratory Animals, 8^th^ edition; and the standards recommended by the United Kingdom’s National Centre for the Replacement, Refinement, and Reduction (NC3R^S^) of Animals in Research: “Responsibility in the Use of Animals in Bioscience Research”. NSG mice (NOD.Cg-*Prkdc^scid^ Il2rg^tm1Wjl^*/SzJ) were bred in-house at The Wistar Institute. Mice were enrolled at 6-8 weeks of age and housed in micro-isolator cages in a designated, specific pathogen-free facility where they were provided with sterile food and water *ad libitum*. Mice were euthanized via CO_2_ administration according to AALAC euthanasia guidelines.

### Tumor Implantation, Grouping, and Equalization

Mice (five males and five females per treatment group) were engrafted subcutaneously with a cell suspension (>98% viability) of 1X10^6^ LCLs (NHC1 LCL-mcherry/enhanced luciferase); these cells were transduced to express mcherry/eLUC using a second generation lentiviral system with pCDH-EF1alpha-eFFly-mCherry (Addgene,Watertown, MA). Cells were implanted subcutaneously using a 27-gauge needle. Animals were weighed three times per week and monitored daily. The Spectrum IVIS CT Bioluminescent Imaging System (Perkin-Elmer; Waltham, MA) was used to randomize mice into groups of 10 per treatment on day five post engraftment to ensure that the average Flux (Photons/sec) was equivalent across groups and to monitor cell growth throughout the study. For imaging studies, Mice were injected with D-luciferin (Gold Biotechnology, St. Louis, MO), i.p. at a dose of 7.5 mg/kg in a dose volume of 10 ml/kg body weight 15 minutes prior to imaging; this was the optimal interval between luciferin injection and bioluminescent imaging as determined by an initial kinetic curve for these cell lines in mice. Mice were anesthetized using isoflurane prior to imaging. Total body Flux (Photons/Sec) was quantitated throughout the study using the Living Image software.

### FK866 Formulation and Treatment Schedule

FK866 was formulated immediately prior to dose. The vehicle was 4% Dimethylacetamide (DMAc), 6% of a 10% Kolliphor solution in DPBS, and 90% DPBS. The vehicle control comprised formulation reagents without compound. FK866 was administered at a dose of 20 mg/kg, i.p.,q.d. in a dose volume of 10 ml/kg body weight.

### Statistical analysis

All experiments presented were conducted at least in duplicate to ensure reproducibility of results. The Prism statistical software package (GraphPad) was used to identify statistically significant differences between experimental conditions and control samples, using statistical tests as indicated in the figure legends.

### Datasets

The data generated in this study are available under the following GEO Accession Number: GSE310487 (RNA-seq mouse), GSE310007 (ATAC-seq), GSE310008 (CUT&Tag), GSE310009 (ChIP-seq), GSE310010 (RNA-seq), GSE310011 (TopoLink), GSE311049 (HiC) and GSE351715 (HiChIP).

Metabolomics data is available on Metabolomics Workbench, Project ID (PR003445; temporarily located at https://dev.metabolomicsworkbench.org:22222/data/DRCCMetadata.php?Mode=Project&ProjectID=PR003445&Access=ExlZ3571

Data shown in Supplemental Figure 1A were obtained from these deposited experiments: GSE170245 (ATAC-seq, ENCODE), GSM935386 (POLR2A ChIP-seq, ENCODE), GSM5360538 (EBNA2 ChIP-seq), GSE49338 (EBNA-LP ChIP-seq), GSM733772 (H3K4me1 ChIP-seq, ENCODE), GSE95899 (H3K4me3 ChIP-seq, ENCODE), GSM733771 (H3K27ac ChIP-seq, ENCODE) and GSE101498 (H3K27ac HiChIP).

The histone proteomics raw files are available online on the repository ProteomeXchange (PRIDE) under the project number PXD069355.

Data for proteomics shown in Figure 1C have been deposited in the ProteomeXchange Consortium through the MassIVE repository under the dataset identifier PXD069549 and MSV000099520.

## Results

**EBV infection triggers NAD production through activation of NAD salvage pathway.**

In our previous studies, we demonstrated that EBV infection activates PARP1, which is essential for driving epigenetic modifications across the viral genome. These PARP1-dependent changes are critical for maintaining viral gene expression and promoting virus-induced cell proliferation^20,75^. Given that sustained PARP1 activity depends on continuous replenishment of NAD^76,77^, we hypothesized that EBV-infected B-cells, in contrast to uninfected cells, upregulate NAD biosynthetic pathways to elevate intracellular NAD levels (**Figure 1A**). To test this hypothesis, we interrogated our previously reported liquid chromatography-mass spectrometry (LC-MS) profiling of polar metabolites in matching B-cells from healthy donors before and after 60 days post-infection with EBV^78^; when infected cells correspond to the lymphoblastoid cell line (LCL) stage. We observed that NAD was one of the top metabolites that significantly increased in B-cells after EBV infection (**Figure 1B**). Consistent with previous report^40,79^, we observed that EBV infection also altered the levels of other metabolites related to specific NAD biosynthetic pathway as in infected cells we observed increased levels of nicotinamide and lower levels of quinolinic acid compared to uninfected B-cells (**Figure 1B**). Nicotinamide is generated during PARP-mediated PARylation and converted into NAD through the salvage pathway (**Figure 1A**). Therefore, our metabolomic analysis indicates that EBV infection promotes NAD biosynthesis predominantly via the NAD salvage pathway. To further support this finding, we performed a global proteomic analysis to determine whether changes in the levels of enzymes involved in NAD metabolism occurred after EBV infection of B-cells. Using mass spectrometry, we compared the protein levels of matching B-cells before and after EBV infection and observed that EBV infection increased the protein levels of NAMPT (**Figure 1C**), the rate-limiting enzyme in the NAD salvage pathway (**Figure 1A**), while reducing the abundance of IDO1 (**Figure 1C**), which catalyzes the initial and rate-limiting step in the kynurenine pathway essential for NAD de novo synthesis (**Figure 1A**).

To examine the contribution of the NAD⁺ salvage pathway to NAD⁺ availability in latently infected cells, we treated two independently generated lymphoblastoid cell lines (LCLs) established from separate donors with FK866{Hasmann, 2003 #329}, a potent and selective inhibitor of NAMPT, the rate-limiting enzyme in this pathway. To assess the on-target effects of FK866, we also treated LCLs with FK866 supplemented with nicotinamide mononucleotide (NMN) or with a NAMPT-targeting PROTAC (**Figure 1D**). NMN bypasses NAMPT inhibition by providing a downstream precursor for NAD⁺ synthesis, whereas the PROTAC provides a complementary approach through NAMPT degradation. A bioluminescent assay showed that both FK866 and the NAMPT-targeting PROTAC reduced cellular NAD⁺ levels, while NMN supplementation restored NAD⁺ levels in FK866-treated cells (Figure 1E).

To further confirm the specificity of NAMPT inhibition by FK866, we performed untargeted global metabolomic profiling of LCLs treated with FK866, FK866 supplemented with nicotinamide mononucleotide (NMN), or a NAMPT-targeting PROTAC (**Figure 1F**). FK866 treatment produced selective metabolic alterations that were largely restricted to metabolites within the NAD⁺ salvage pathway, consistent with on-target inhibition of NAMPT rather than widespread metabolic disruption. Supplementation with NMN restored these metabolic changes (**Figure 1F**), demonstrating that the observed effects resulted from depletion of the NAMPT-dependent NAD⁺ salvage pathway and could be rescued by bypassing the enzymatic block. Furthermore, degradation of NAMPT using the PROTAC induced metabolomic changes that closely mirrored those observed following FK866 pharmacological inhibition (**Figure 1F**), providing independent evidence that the metabolic phenotype is a direct consequence of NAMPT inhibition.

Finally, the effects of FK866 treatment on cell viability and proliferation were assessed. We found that treatment with 2.5 nM FK866 had no significant effects on cell viability at 3 days and no detectable effect on cell proliferation . However, at higher concentrations FK866 treatment resulted in increased toxicity and cell death (**Supplemental Figure 1**). Therefore, 2.5 nM FK866 was selected for all subsequent experiments, allowing interrogation of NAD salvage pathway inhibition while minimizing confounding effects on cell viability and growth. Together, these findings validate the specificity of FK866, confirm that its metabolic effects are mediated through disruption of NAD⁺ salvage, and demonstrate that EBV infection rewires cellular NAD metabolism to favor the salvage pathway.

### NAMPT activity is essential during primary EBV infection of B-cells

A recent study demonstrated that IDO1 plays a critical role during the early phase of EBV infection by supporting NAD production^40^. We therefore aimed to determine whether the NAD salvage pathway similarly contributes to NAD biosynthesis in the initial stages of EBV infection. Analysis of publicly available proteomic datasets from a time-resolved experiment evaluating global protein abundance in B-cells over a 28-day period following EBV infection^80^ showed that NAMPT levels increase as early as one day post infection and reaching peak abundance and stabilizing by day four. Notably, the increase in NAMPT expression closely followed peak expression of the viral transcriptional regulators EBNA2 and EBNA-LP, suggesting a potential correlation between the expression of latent viral proteins and NAMPT expression (**Supplemental Figure 2A**).

We observed that the increase in NAMPT protein abundance correlates to an observed increase in NAD levels in the 48 hours following EBV infection when measured using a luminescent assay (**Figure 1G**). Since EBV infection induces CD23 expression in B-cells^81,82^, we used CD23 positivity as a marker for successful viral infection of primary B-cells. We infected primary B-cells with EBV and after 48 hours we sorted them for CD23 expression and measured NAD levels. We observed that compared to uninfected cells (time 0) EBV infected B-cells (CD23+) had significantly higher NAD levels (**Figure 1G**). Additionally, we infected B-cells from a separate donor and treated the cells with 2.5nM of FK866 at the time of infection and observed a significant decrease in NAD levels two days post infection (**Figure 1H**), confirming that the increased NAD availability was due to activation of the salvage pathway.

Next, to confirm a positive correlation between NAD pathway activation and the extent of EBV infection, we tested whether depletion of NAD levels by the salvage pathway blockade affected B cells response to EBV infection. Using flow cytometric assay to quantify CD23+ expression as a proxy of EBV infection, we observed that FK866 treatment significantly reduced the percentage of B-cells infected by the virus (**Figure 1I**). To confirm the on-target effect of FK866-mediated NAMPT inhibition, we supplemented cells with NMN (**Figure 1I**). Supplementation with NMN restored EBV infection levels in B-cells treated with FK866. Further supporting these results, microscopic evaluation demonstrated that FK866 treatment markedly decreased the number and size of proliferating B cell clusters typically observed following infection of primary B cells with EBV (**Figure 1J**). NMN supplementation reversed these effects, restoring both cluster formation and growth (**Figure 1J**). These observations suggest that NAD salvage is critical during the early stages of EBV infection of B cells.

### EBV influences NAMPT expression by altering the chromatin structure of its promoter

The EBV protein EBNA2 regulates viral and host gene expression and is essential for EBV-mediated B-cell immortalization^27,31,32,83,84^. Similarly, EBNA-LP plays a critical role in modulating host gene expression^33,34,85–89^. We and others recently reported that EBNA-LP influences host transcription through the regulation of chromatin looping, thereby altering the three-dimensional structure of the genome^35,90^. To explore a potential role for EBNA2 and EBNA-LP in regulating NAMPT expression, we analyzed publicly available ChIP-seq datasets for EBNA2 and EBNA-LP binding across the NAMPT genomic locus in EBV-positive LCLs^91,92^ as well as publicly available ENCODE datasets on GM12878 LCLs, including RNA Pol II ChIP-seq to mark active promoters, ATAC-seq to identify regions of open chromatin, and H3K27ac HiChIP to detect chromatin interactions connecting regulatory regions. We observed strong RNA Pol II binding and prominent open chromatin signals (ATAC-seq peaks) at the NAMPT promoter, confirming its active transcriptional state in EBV-infected LCLs (**Supplemental Figure 2**). EBNA-LP was found to occupy the NAMPT promoter and several upstream regions, including a notable site located approximately 200 kb upstream (**Supplemental Figure 2B**). Remarkably, this distal region was the only site where EBNA2 binding was detected. Analysis of H3K27ac HiChIP data^93^ revealed that the distal locus co-occupied by EBNA2 and EBNA-LP physically interacts with the NAMPT promoter (**Supplemental Figure 2**). These findings show that EBNA2 and EBNA-LP occupy regions that may contribute to NAMPT regulation.

To determine whether the candidate enhancer occupied by EBNA2 and EBNA-LP contributes to NAMPT expression, we targeted this region using dCas9–KRAB-mediated CRISPR interference^94^ with two independent sgRNAs (sg1 and sg2). Controls included a scrambled sgRNA and an sgRNA targeting a nearby region outside the predicted enhancer (sg3) (**Supplemental Figure 2C**). Targeting the candidate enhancer reduced NAMPT protein levels relative to both controls, whereas targeting the adjacent region had no effect (**Supplemental Figure 2D**). These results demonstrate a functional role for the EBNA2- and EBNA-LP-bound regulatory region in maintaining NAMPT expression in EBV-transformed cells.

### NAD inhibition affects transcriptional activation of B-cells by EBV

To determine whether the NAD salvage pathway is necessary for EBV-mediated transcriptional reprogramming of the B-cells, we performed global transcriptomic analysis of LCLs treated with a sub-lethal dose of the NAMPT inhibitor FK866^95^. Principal component analysis (PCA) revealed that NAMPT inhibition accounted for 65% of the variance on transcriptomic profiles between FK866-treated and untreated cells (**Figure 2A**).

**Figure 2:**
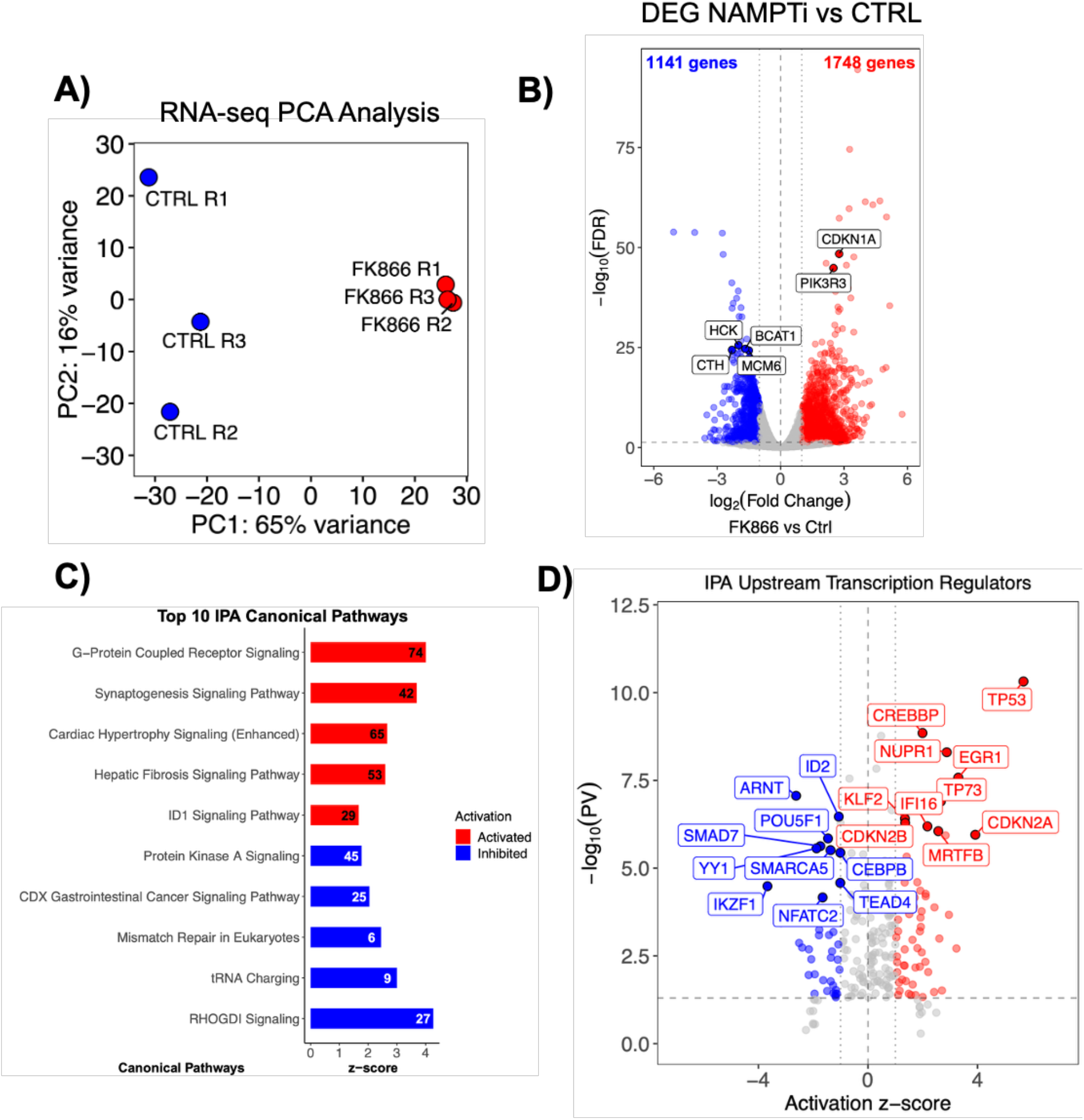
NAMPT inhibition reprograms cellular signaling and cytoskeletal remodeling and prevents plasma cell differentiation. RNAseq results from three technical replicates of LCLs treated with 2.5nM FK866 for three days. Principal Component Analysis (PCA) based on RNAseq shows FK866 treatment is responsible for the majority of differences in gene expression between control and treated conditions (**A**). Volcano plot showing differentially expressed genes (blue = downregulated, red = upregulated, grey = nonsignificant) (**B**). Bar plot showing the top 10 activated/inhibited canonical pathways identified by Ingenuity Pathway Analysis (IPA) (**C**). Volcano plot displaying the up- (red) and downregulated (blue) upstream transcription regulators identified by IPA based on differential gene expression (**D**).

NAD salvage pathway inhibition resulted in widespread transcriptional changes, with both upregulation and downregulation of host genes (**Figure 2B**). In total, 2,889 genes were significantly differentially expressed (FDR < 0.05, |log2 fold change| ≥ 1), including 1,748 upregulated and 1,141 downregulated transcripts in FK866-treated cells (**Figure 2B**). Notably, several genes involved in proliferation and B cell function — such as *PIK3R3*, and *CDKN1A* (*p21*) — were among the most significantly upregulated (**Figure 2B**). In addition, key genes related to cellular metabolism (*CTH*, *BCAT1*) and B-cell receptor (BCR) signaling and proliferation (*HCK*, *MCM6*) were markedly reduced (**Figure 2B**). Pathway enrichment analysis indicated that inhibiting NAMPT in B-cells resulted in a unique transcriptional and signaling profile. This profile was characterized by the activation of G-protein-coupled receptor (GPCR) signaling, synaptogenesis, and ID1 signaling (**Figure 2C**). Additionally, there was an inhibition of protein kinase A (PKA) pathways, mismatch repair mechanisms, and RhoGDI pathways (**Figure 2C**). Together, these analyses show that NAMPT inhibition alters the expression of genes involved in metabolism, cell-cycle regulation, and DNA repair in EBV-transformed LCLs.

The upstream transcription regulators analysis of our RNA-seq data suggested a broad host stress and checkpoint response counteracting EBV-driven proliferation in B-cells. Using stringent analysis conditions (|z-score| >= 2, q<0.01) upstream prediction identified activation of TP53, TP73, CDKN2A/B, CREBBP, consistent with a transcriptional response associated with tumor-suppressive checkpoints, and activation of KLF2, a key regulator of B-cell quiescence and a barrier to plasma cell differentiation^96,97^ (**Figure 2D**). Predicted activation of IFI16 and NUPR1, EGR1 also suggested antiviral sensing and stress-adaptive signaling^98–100^ being induced by NAD depletion (**Figure 2D**). Interestingly, the histone acetyltransferase CREBBP (CBP) was also predicted to be activated by FK866 treatment (**Figure 2D**), suggesting induction of chromatin remodeling programs in response to NAD salvage inhibition.

In contrast, TFs that normally sustain B-cell developmental programs and inflammation like IKZF1, ID2, NFATC2, and CEBPB^101–104^ were predicted to be inhibited. Consistent with a possible reduction in the plasticity of transcriptional programs governing B-cell differentiation and function, the pluripotency factor POU5F1 (OCT4) was also predicted to be inhibited following NAD⁺ depletion (**Figure 2D**). Loss of metabolic adaptation was also suggested by prediction of ARNT inhibitions in FK866 treated cells (**Figure 2D**). In addition, TFs involved in chromatin remodeling were predicted to be inhibited including SMARCA5, and YY1, which has been recently identified to interact with EBNA-LP and participate in remodeling of the three-dimensional structure of both host and viral genome^35,85,90^ (**Figure 2D**).

Collectively, these results indicate that NAMPT inhibition is associated with extensive transcriptional reprogramming, including changes in genes and predicted regulators linked to proliferation, cellular stress, antiviral responses, and chromatin organization. This pattern is consistent with a shift away from the proliferative transcriptional state associated with EBV latency toward a stress-responsive state.

### Chromatin remodeling in EBV-infected cells requires NAD salvage pathway activation

To determine whether the transcriptional changes induced by NAD salvage pathway inhibition are accompanied by chromatin remodeling, as suggested by upstream regulator analysis, we performed ATAC-seq on LCLs before and after FK866 treatment. Principal component analysis (PCA) showed that FK866 treatment accounted for 24% of the variance in chromatin accessibility between treated and untreated cells (**Figure 3A**). ATAC-seq profiling revealed changes in the distribution of accessible chromatin in B-cells following NAD depletion (**Figure 3B**). Quantitative proteomic analysis of histone modifications revealed no major global alterations in marks associated with open chromatin including H3K4me1, H3K4me3, and H3K9ac. NAMPT inhibition reduced H3K27ac levels by approximately 38%, while the abundance of the other measured histone modifications remained largely unchanged (**Supplemental Figures 3A** and **3B**). Despite this reduction in total abundance, the genomic pattern of H3K27ac enrichment was largely preserved following NAD⁺ depletion (**Supplemental Figure 3C**). Thus, the observed changes in chromatin accessibility occurred alongside limited changes in the distribution of H3K27ac.

**Figure 3:**
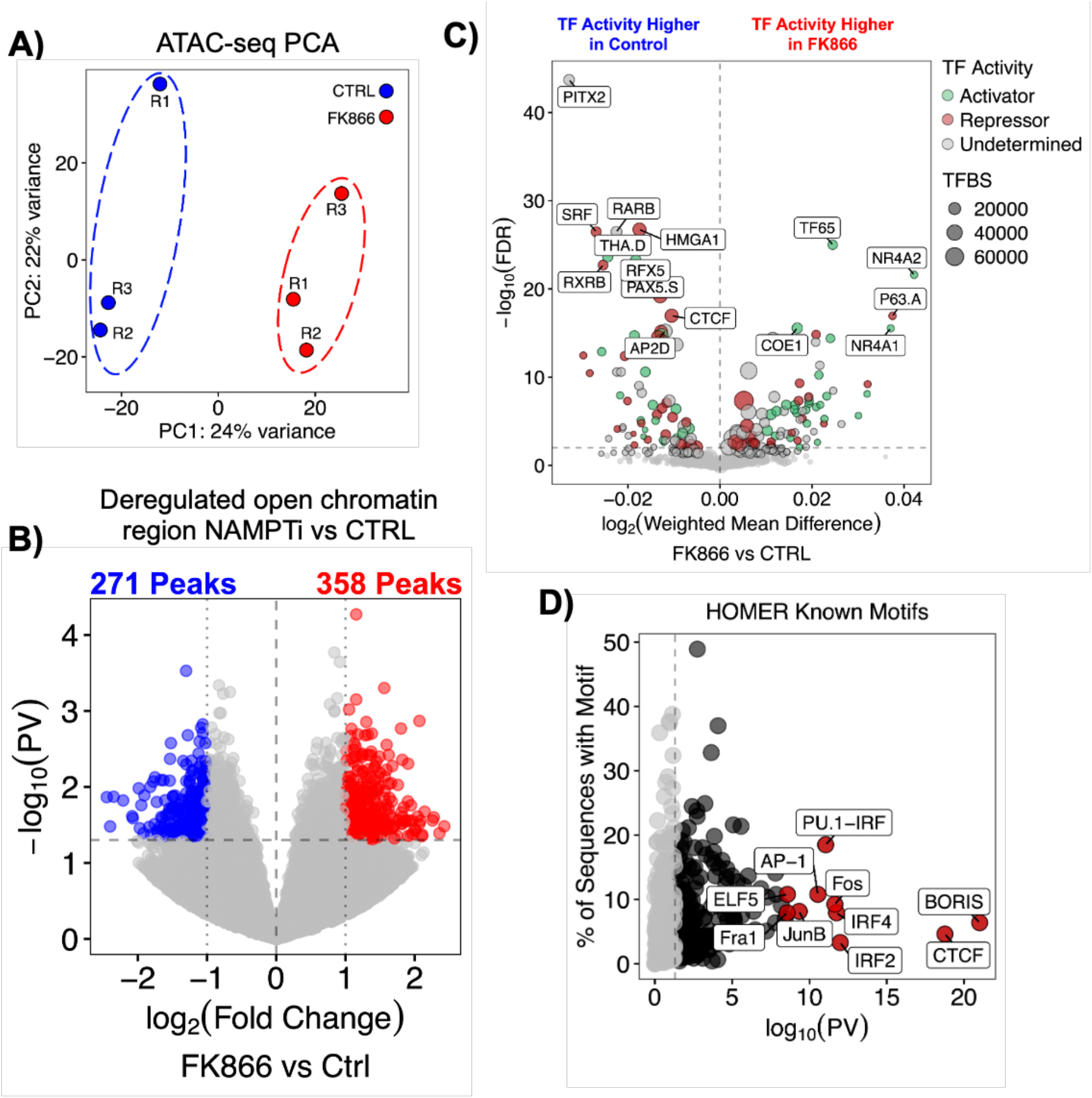
NAMPTi induces changes in chromatin accessibility of CTCF/cohesin binding sites: ATACseq results from three technical replicates of LCLs treated with 2.5nM FK866 for three days. Principal Component Analysis (PCA) based on ATACseq counts (**A**). Volcano plot showing differentially accessible chromatin regions (blue = closed, red = open, grey = nonsignificant) (**B**). Volcano plot showing transcription factor motifs associated to differentially accessible chromatin regions from ATACseq. The colors of the dots represent whether the peak is associated to an up- (green) or downregulated (red) gene according to RNAseq under the same conditions, while the size of the dots are proportional to the number of motifs found that correspond to the indicated transcription factor (**C**). Scatterplot showing enriched TF motifs identified by HOMER findMotifsGenome function in the differentially accessible regions. Red dots are the top 10 enriched motifs (**D**).

Overall, NAMPT inhibition altered chromatin accessibility, with both gains and losses in accessible regions (**Figure 3B**). In total, 629 regions exhibited significant changes in chromatin accessibility (p-value < 0.05, |log₂ Fold Change| > 1), including 358 regions with increased accessibility (**Figure 3B**, red circles) and 271 regions with decreased accessibility (**Figure 3B**, blue circles). In untreated control cells, accessible chromatin regions were enriched for motifs recognized by B-cell lineage-defining and chromatin architectural transcription factors — such as PAX5, POU2F1/2 and CTCF — as well as regulators of metabolic and cytokine responses including SMAD4 and KLF12 (**Figure 3C**). Following FK866 treatment, this motif landscape was substantially altered. Newly accessible regions were enriched for binding motifs of NF-κB subunit REL, RUNX3, NR4A1 and TP63 (**Figure 3C**), suggesting a shift toward transcriptional programs associated with immune activation and stress responses.

Notably, motif analysis identified the CTCF binding motif among the most significantly enriched motifs in regions with altered chromatin accessibility following FK866 treatment. Given the role of CTCF in higher-order chromatin organization, this finding raises the possibility that changes in CTCF-associated regulatory regions contribute to the chromatin alterations observed following NAMPT inhibition. Taken together, these motif changes indicate that inhibition of the NAD⁺ salvage pathway alters the chromatin accessibility landscape and suggest potential changes in the transcriptional regulatory programs associated with immune and stress responses.

### Cohesin-CTCF chromatin loop extrusion requires activation of the NAD salvage pathway

In our previous work, we demonstrated that CTCF and cohesin binding across the genome in EBV-infected cells requires PARP1 activity^18–20^. Importantly, others have reported that EBV-transformed cells are highly dependent on PARP1 for DNA repair^105^, highlighting its involvement in multiple chromatin-associated processes. Because PARP1 requires NAD⁺ as a substrate, NAD⁺ depletion could limit its activity and thereby affect the binding of architectural proteins. We therefore hypothesized that NAD⁺ depletion would alter CTCF and cohesin occupancy, with reduced PARP1 activity as a potential mediator. Consistent with this model, motif analysis of accessible chromatin regions, showed a significant enrichment for CTCF binding motifs among regions with altered chromatin accessibility following NAD depletion (**Figure 3D**), suggesting that regions associated with

CTCF-mediated chromatin organization are affected by NAMPT inhibition. Thus, we determined the impact of NAD depletion on higher-order chromatin organization. To directly assess this, first we examined CTCF and cohesin occupancy in EBV+ LCLs upon FK866-mediated NAMPT inhibition. CTCF and cohesin interaction is essential for chromatin loop formation^4,106^, which is initiated by NIPBL binding to DNA, recruiting and loading the cohesin complex—comprising SMC1, SMC3, and RAD21^107,108^. Once loaded, cohesin complex translocates along the DNA until it encounters convergent CTCF-binding sites, where it stabilizes loop formation^109^ (**Figure 4A**). To examine the effect of NAD salvage pathway inhibition on this process, we first confirmed that protein level of SMC3 and CTCF remains stable after treatment with 2.5nM FK866 for three days does (**Figure 4B**). Next, we performed ChIP-seq for CTCF, the cohesin subunit SMC3, and the cohesin loader NIPBL in LCLs before and after FK866 treatment (**Figure 4C**). Heatmaps of normalized ChIP-seq signal (z-scored fragments per kilobase per million reads), hierarchically clustered by signal intensity, revealed three major clusters with distinct occupancy profiles (**Figure 4C**). Cluster 1 comprised regions that showed increased SMC3 and NIPBL occupancy but minimal CTCF binding after FK866 treatment, suggesting altered cohesin occupancy at regions associated with cohesin loading (**Figure 4C**). Cluster 2 represented regions with the strongest CTCF, SMC3, and NIPBL occupancy before treatment, where all three signals increased further following NAD⁺ depletion (**Figure 4C**). This pattern is consistent with parallel increases in CTCF, cohesin, and cohesin-loader occupancy at pre-existing sites. Cluster 3 consisted of regions with CTCF enrichment greater than that observed in cluster 1 but largely unchanged (or modestly increased) following FK866 treatment. SMC3 and NIPBL were also present at these sites before treatment, and their occupancy increased following NAD⁺ depletion (**Figure 4C**). This pattern is consistent with increased cohesin recruitment or retention at pre-existing CTCF-bound sites.

**Figure 4.**
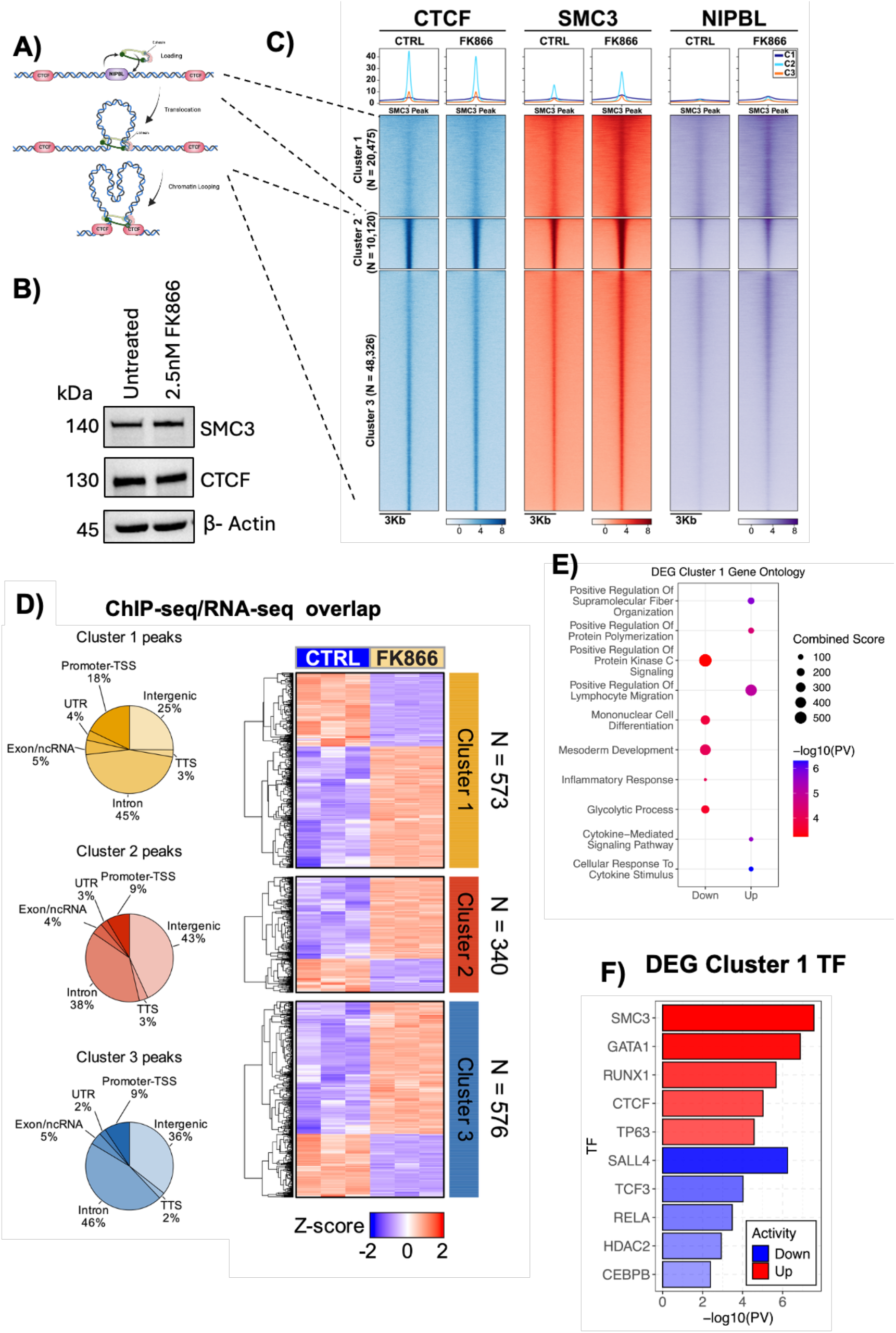
**FK866 treatment of LCLs induces genome-wide retention of cohesin complex on chromatin**: Model of how cohesin complex is loaded onto chromatin and how it extrudes loops (**A**). Heatmaps displaying ChIP-seq signal for CTCF (blue), SMC3 (red) and NIPBL (purple) genome-wide before and after NAMPT inhibition. Heatmaps were divided into three clusters using unsupervised clustering based on ChIP-seq signal. Peaks belonging to each clusters are indicated on the side Western blot showing that FK866 treatment does not alter the protein levels of CTCF and SMC3 (**B**). **(C**). Pie chart of annotated peaks in each cluster in B indicating the localization of peaks at promoter-TSS, intergenic, exon/ncRNA, intron, UTR and transcription termination site (TTS) gene regions (left) and heatmap showing Z-scores of overlapping differentially expressed genes from the RNA-seq experiment (right) (**D**). Dot plot of enriched Gene Ontology pathways based on DEGs overlapping with cluster 1 peaks (**E**). Bar plot showing enriched transcription factors based on ENCODE ChIP-seq experiments on up- (red) and downregulated (blue) genes (**F**).

To further characterize regions with altered cohesin occupancy in all three clusters we first annotated their genomic distribution. In cluster 1, approximately 25% of peaks mapped to intergenic regions and 18% to promoters and transcription start sites (TSS). Clusters 2 and 3 showed a greater proportion of intergenic peaks, accounting for 43% and 36%, respectively, whereas promoter/TSS-associated peaks accounted for 9% in each cluster (Figure 4D, left). The proportion of promoter-associated sites in cluster 1 was therefore approximately twice that in clusters 2 and 3.

We next compared genes associated with each cluster with our RNA-seq dataset (**Figure 4D**, right). Following FK866 treatment, we identified 573 differentially expressed genes in cluster 1 (357 upregulated and 216 downregulated), 340 in cluster 2 (241 upregulated and 99 downregulated), and 576 in cluster 3 (390 upregulated and 186 downregulated). These represented approximately 7%, 4%, and 3% of the genes associated with each cluster, respectively. The higher proportion of differentially expressed genes in cluster 1 coincided with its greater representation at promoters. Among genes showing significant expression changes, the response at sites of increased cohesin occupancy was skewed toward upregulation across all three clusters. Most associated genes, however, showed no significant change in expression indicating that changes in CTCF, SMC3 and NIBPL observed were largely independent of changes in transcription of underlying loci.

Because cluster 1 contained the highest proportion of differentially expressed genes, we performed Gene Ontology analysis of these genes to identify the biological processes associated with their expression changes. Gene ontology analysis of the target genes associated to cluster 1 using *enrichR* software^110^ revealed enrichment for pathways associated with cytokine production, inflammatory signaling, and cellular activation, suggesting that the transcriptional changes associated with altered cohesin occupancy involve immunoregulatory programs in B-cells (**Figure 4E**). Analysis of transcription factors associated with deregulated genes within cluster 1 showed enrichment of CTCF and SMC3 binding sites, suggesting preferential activation of genes positioned within chromatin loops, together with motifs for RUNX1, GATA1, and TP63, consistent with re-engagement of hematopoietic differentiation programs and p53-family stress responses (**Figure 4F**).

CTCF was implicated in both the chromatin accessibility and transcriptional analyses. CTCF binding motifs were enriched in accessible regions in untreated cells (**Figure 3C**), while gene-expression changes in cluster 1 predicted increased CTCF regulatory activity following FK866 treatment (**Figure 4F**). At these cluster 1 sites, however, CTCF occupancy remained relatively low and changed only modestly, whereas SMC3 and NIPBL occupancy increased. The CTCF-associated transcriptional signature therefore coincided with increased cohesin and NIPBL occupancy rather than a comparable increase in CTCF binding.

Together, these findings demonstrate that inhibition of the NAD⁺ salvage pathway alters CTCF, cohesin, and NIPBL occupancy across the genome in EBV-infected B cells, alongside changes in chromatin accessibility and gene expression. These results support a role for NAD⁺ salvage pathway activity in maintaining chromatin organization and transcriptional regulation in EBV-transformed cells.

### NAD⁺ salvage inhibition is associated with changes in pre-existing chromatin contacts

Our ChIP-seq data indicated that NAD⁺ salvage pathway activity contributes to the regulation of CTCF and cohesin occupancy in EBV-infected cells. To examine whether these changes were accompanied by alterations in three-dimensional genome organization, we performed Hi-C in control and FK866-treated LCLs. Principal component analysis (PCA) separated the two conditions along a component accounting for 65% of the variance in the analyzed Hi-C data (**Figure 5A**). We first examined chromatin organization at the compartment level. A compartments are generally enriched in gene-rich, transcriptionally active regions, whereas B compartments are associated with gene-poor, transcriptionally repressed regions^111^ (**Figure 5B**). When we interrogated our HiC data to assess A and B compartment before and after FK866 treatment we observed changes in the spatial organization of chromatin, where 707 genomic regions transition between A (active) and B (inactive) compartments, including 492 compartments that switch from A to B (∼70% of transitioning compartments), and 215 that switch from B to A (∼30% of transitioning compartments) (**Figure 5C**). This data suggests that NAD⁺ salvage pathway inhibition alters large-scale chromatin compartmentalization, alongside the changes in chromatin accessibility observed by ATAC-seq (**Figure 3B**). We then tested whether A/B compartment transitions induced by FK866 treatment were accompanied by corresponding transcriptional changes. We compared expression changes of genes located in switched versus stable compartments. The log2 Fold Change distributions of genes undergoing A-to-B (n = 35) and B-to-A (n = 63) transitions largely overlapped with those of genes in stable A (n = 9982) or stable B (n = 1321) compartments (**Supplemental Figure 4A**). Relative to stable genes of the same compartment of origin, we detected no significant location shift in log2 Fold Change for either A-to-B genes (estimated shift ∼ 0.06, 95% CI −0.16 to 0.29; p = 0.59) or B-to-A genes (∼ −0.07, 95% CI −0.23 to 0.09; p = 0.38) (**Supplemental Figure 4B**). Consistent with this, the magnitude of compartment change (ΔE1) was only weakly associated with mean log2 Fold Change per bin, explaining 0.5% of its variance (R^2^ = 0.005) (**Supplemental Figure 4C**). These data together indicate that compartment changes following FK866 treatment are largely uncoupled from steady-state transcriptional output.

**Figure 5.**
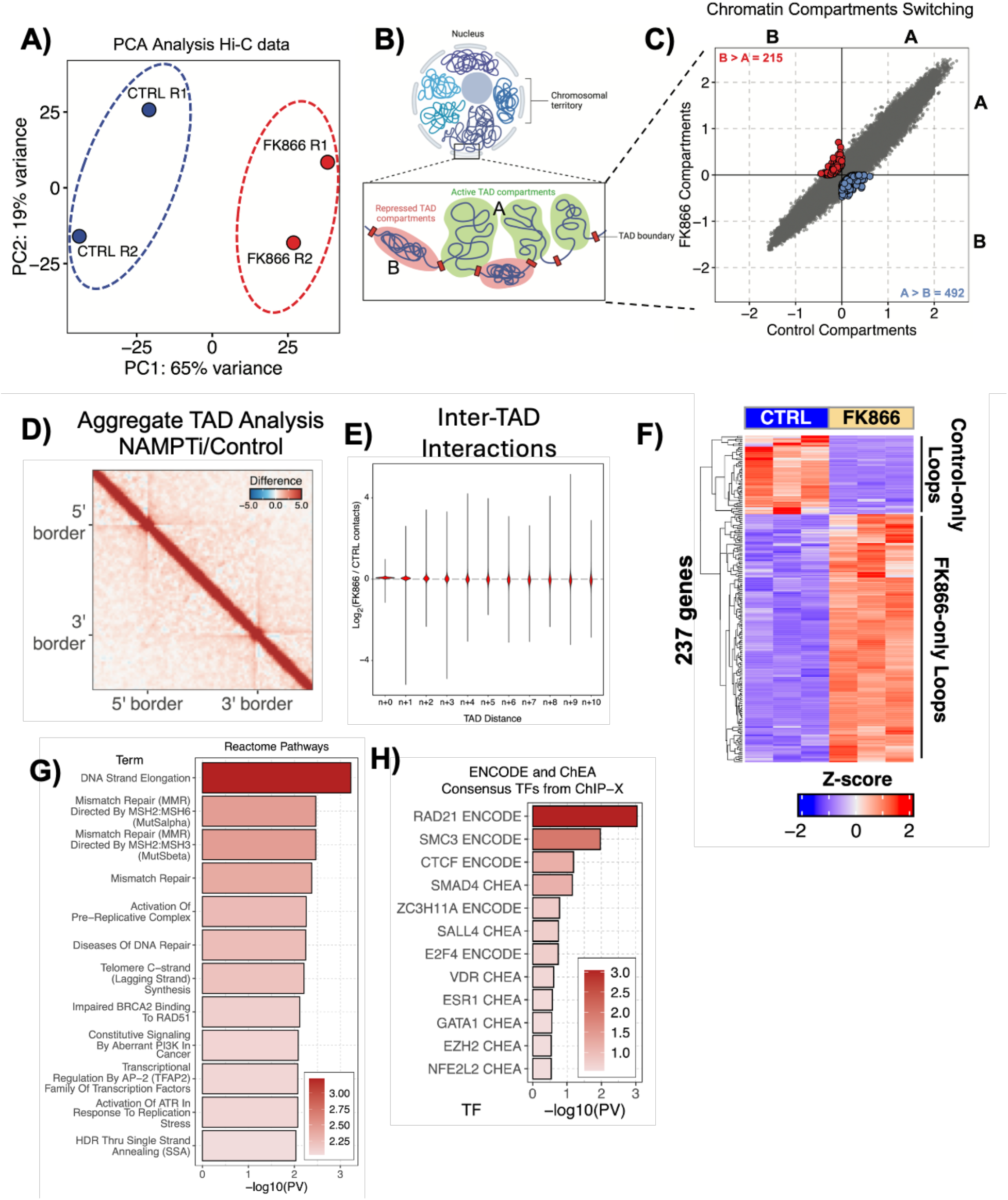
**FK866 treatment leads to global changes in chromatin architecture**: PCA shows FK866 treatment induces changes in chromatin architecture from two biological replicates (**A**). Model of how chromatin is organized in A (active) and B (inactive) compartments inside the nucleus (**B**). Scatterplot showing A and B compartment detected by HiC assay upon NAMPTi in LCLs. Compartment are detected using 100kb bins and those that switch from A to B and vice versa are colored in blue and red respectively (P-value < 0.05) (**C**). Aggregate TAD Analysis (ATA) showing the difference between FK866-treated and Control matrices (**D**). Violin plots showing the log_2_ difference between inter-TAD interaction counts between FK866-treated and Control matrices (**E**). Heatmap showing the Z-score of differentially expressed genes that overlap with either control-only and FK866-only loop anchors detected in the HiC experiment. Only concordant genes (log_2_FC > 1 & FK866-only loop or log_2_FC < −1 & control-only loop overlaps) are reported (**F**). Barplot displaying the Reactome Pathways identified by enrichR pathway analysis based on concordant genes showed in the heatmap in (F) (**G**). Barplot showing ENCODE TFs enriched at overlapping genes in (F) (**H**).

To examine the effects of NAD⁺ depletion at a finer scale, we next analyzed topologically associating domains (TADs), which are contiguous genomic regions within compartments that are characterized by more frequent contacts within each domain than between neighboring domains (**Figure 5B**). We interrogated our HiC dataset to assess the FK866 effect on TADs. The Aggregate TAD Analysis (ATA) revealed that FK866 treatment was associated with increased definition of TAD boundaries, as evidenced by sharper domain borders (**Figure 5D**). Analysis of inter-TAD interactions also revealed a reduction in long-distance contacts between TADs separated by seven or more domains (n+7 and higher) following NAD⁺ salvage pathway inhibition (**Figure 5E**). These findings are consistent with reinforcement of domain organization, with greater insulation of individual domains and reduced long-range interdomain contacts following NAD⁺ depletion.

Next we examined chromatin loops identified from the HiC dataset. Of all loops detected (10kb bin size, FDR < 0.01 and present in both replicates), almost 50% were specific to FK866-treated LCLs while around 46% were specific to the control condition. Only a small portion (∼6%) was retained after NAMPT inhibition (**Supplemental Figure 5**). In addition, aggregate peak analysis of regions of altered cohesin occupancy in cluster 1 (**Supplemental Figure 6A**) showed higher frequency of chromatin looping after FK866 treatment (**Supplemental Figure 6B**). Together, these results suggest that NAD⁺ availability contributes to the organization of long-range chromatin contacts.

To investigate the relationship between chromatin looping and gene expression we intersected chromatin loop frequency with our RNA-seq dataset and identified 237 genes exhibiting significant transcriptional changes concordant with loop frequency changes following FK866 treatment (**Figure 5F**). Notably, chromatin interactions with higher contact frequencies in FK866-treated cells than in controls were predominantly associated with transcriptional upregulation, whereas those with lower contact frequencies were associated with transcriptional downregulation (**Figure 5F).** It is worth noting that ∼37% of all overlapping genes identified belong to ChIP-seq cluster 1 (n = 88) (**Supplemental Figure 6C**). These observations suggest a potential association among increased cohesin occupancy, reinforcement of existing chromatin architecture, and changes in gene expression following NAMPT inhibition. The association between chromatin contact frequencies and gene expression prompted us to examine whether the genes linked to these changes shared common biological functions. We therefore performed Gene Ontology analysis to identify cellular processes enriched among the associated genes. DNA strand elongation, mismatch repair, and transcriptional regulation were among the most enriched categories (**Figure 5G**). These findings indicate that the chromatin and transcriptional changes observed following NAMPT inhibition were associated with genes involved in DNA replication and repair, as well as the regulation of gene expression.

To identify regulators associated with genes showing both altered chromatin contacts and differential expression, we performed transcription factor and chromatin regulator enrichment analysis. Enrichment analysis for transcription factors showed that regions of altered chromatin looping were mostly enriched for cohesin [RAD21 and SMC3]/CTCF complex signature, consistent with a role of this factor in regulating chromatin architecture (**Figure 5H**). The analysis also identified regulatory signatures associated with development and stemness programs (SALL4, SMAD4, EZH2, ESR1, VDR), cell cycle and proliferation control (E2F4), response to oxidative stress and inflammation (NFE2L2 and VDR) and differentiation (GATA1 and ESR1). Together, these findings indicate that inhibition of NAD⁺ salvage pathway activity is associated with changes in CTCF/cohesin occupancy, reinforcement of chromatin architecture, and transcriptional programs in EBV-transformed B cells.

### NAD salvage activity influences asymmetric loop extrusion and conformation of highly interactive regulatory elements

In addition to influencing TAD organization, in our Hi-C contact maps we observed that FK866 treatment was associated with changes in chromatin stripes, which represent regions where a single genomic locus interacts with an extended linear stretch of chromatin (**Supplemental Figure 7A**). These structures are indicative of asymmetric loop extrusion and often reflect directional interactions associated with enhancer activity, promoter tethering, or boundary insulation^4,5,112–115^. At our Hi-C resolution (50-100 kb), we were unable to properly appreciate stripes formation, therefore we decided to perform the newly developed TopoLink assay (Dovetail Genomics). This assay leverages the Tn5 ability to cut chromatin at accessible sites making it possible to generate high resolution contact maps while keeping the number of cells to a minimum of 1×10^6^ per condition. By applying this assay, we were able to identify a total of 1277 stripes. Only ∼12% of these were altered in LCLs following FK866 treatment (**Supplemental Figure 7B**, right). Interestingly, while 52 stripes were lost upon NAD salvage inhibition, a significantly larger number — 100 stripes (∼8% of all observed) — emerged de novo in FK866-treated cells (**Supplemental Figure 7B**). Newly formed stripes were frequently located at genomic regions marked by strong CTCF binding and increased cohesin retention, as evidenced by elevated SMC3 and NIPBL ChIP-seq signal compared to untreated controls (**Supplemental Figure 7C**). CTCF chromatin looping is regulated by PARP activity, which requires continuous replenishment of NAD through the salvage pathway (**Supplemental Figure 8A**). Indeed, in LCLs, inhibition of NAD salvage by FK866 disrupts PARP activity by blocking the recycling of nicotinamide into NAD (**Supplemental Figure 8B**). Thus, based on this observation, we expected PARP inhibition to phenocopy NAMPT inhibition. Consistent with our hypothesis, treatment of LCL cells with the PARP inhibitor BMN673 resulted in a similar reduction of stripe formation compared to control cells (**Supplemental Figures 7B** and **7C**). Overall, the observed changes in stripes support a role for NAD levels in influencing the asymmetric loop extrusion model: NAD depletion leads to changes in occupancy of cohesin at one CTCF-bound site, while the opposing end of the loop continues to extrude, generating the characteristic conformational pattern.

### EBV epigenetic landscape responds to NAD changes

We and other groups have previously demonstrated that CTCF and cohesin binding across the EBV genome regulate EBV genome 3D architecture and proper 3D folding is important for EBV latent infection maintenance. Since we observed that NAD levels affect host chromatin architecture, we hypothesized that FK866 treatment could alter EBV chromatin architecture by impairing CTCF and cohesin function. To test this, we employed our EBV-enriched Hi-C approach, as previously described^18,19^, to profile the 3D structure of the EBV genome, and integrated these data with ChIP-seq datasets for NIPBL, SMC3, and CTCF aligned to the viral genome. Consistent with our observations in the host genome, FK866 treatment resulted in pronounced changes in EBV chromatin looping, with both the gain and loss of specific loops across the viral episome (**Figure 6A**). Notably, NAD depletion was associated with an increase in short-range loops, particularly in a region encompassing the origin of lytic replication (OriLyt-Left), the latent promoter Qp, and the lytic promoter Zp, which regulates the lytic transactivator gene *BZLF1* (**Figure 6A**). In addition, we observed that the promoter regions for *LMP1* and *LMP2* established new interactions with the terminal repeat (TR) region and with the locus encoding the small non-coding EBER RNAs (**Figure 6A**). Conversely, several loops were lost following FK866 treatment, including interactions between the Qp and Cp promoters, as well as between Qp and OriLyt-Left (**Figure 6A**). Altered viral chromatin loops were predominantly observed between regions occupied by CTCF, SMC3, and NIPBL (**Figure 6A**). Following FK866 treatment, SMC3 occupancy increased across the EBV genome, whereas CTCF binding was reduced at many viral regions (**Figures 6A and 6B**). This pattern differed from our observations in the host genome, where CTCF occupancy was maintained or increased at specific regions following NAD⁺ depletion. Overall, this shift in chromatin architecture — characterized by enhanced cohesin retention and reduced CTCF occupancy — closely mirrors our previous findings in EBV-infected cells treated with the PARP inhibitor BMN673. These findings are consistent with a model in which NAD availability supports PARP1-dependent regulation of CTCF and cohesion occupancy across the viral episome.

**Figure 6.**
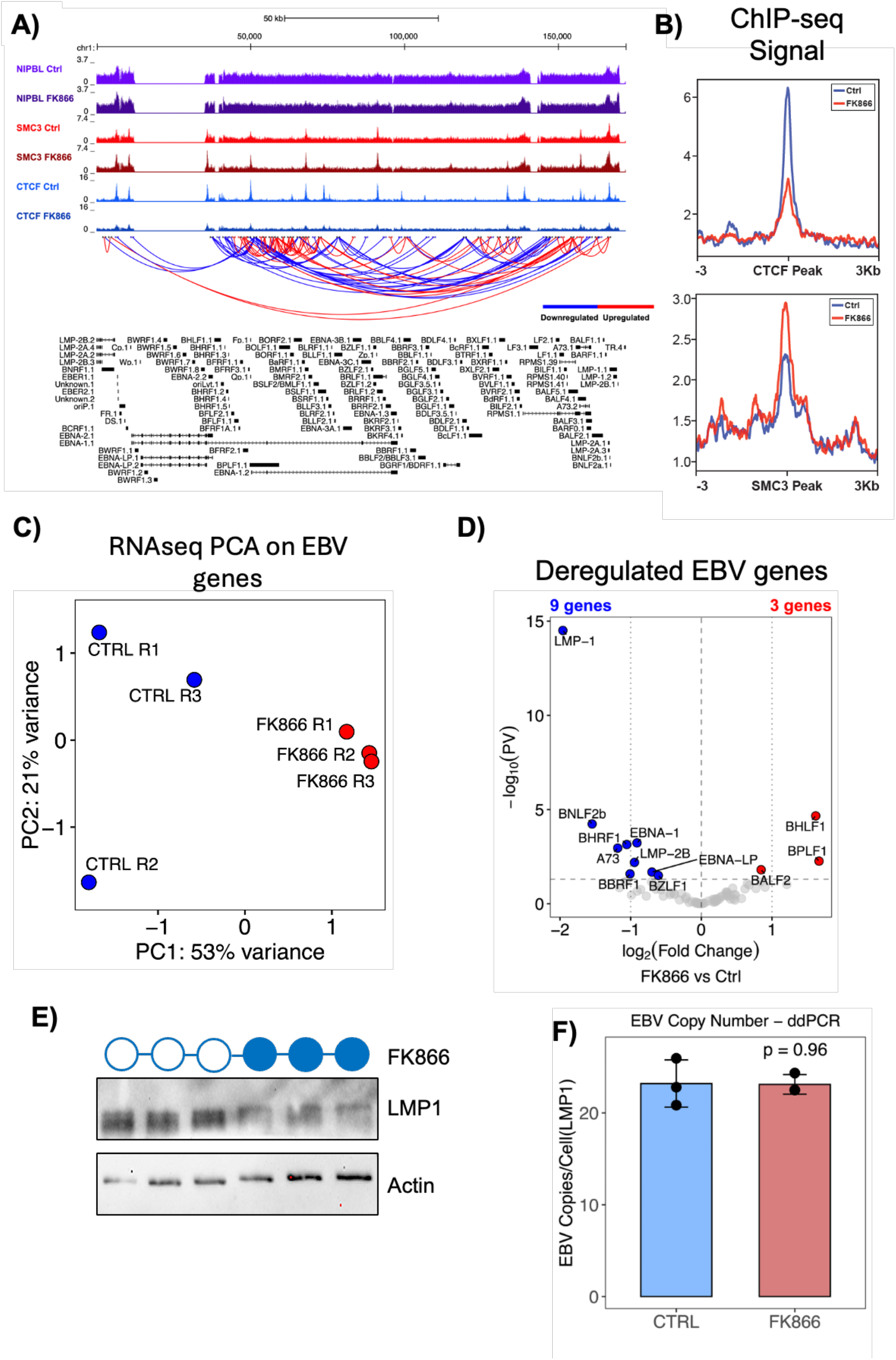
**NAMPT inhibition induces changes in EBV chromatin architecture, CTCF/cohesin complex binding and latent gene expression**: UCSC Genome Browser session showing CTCF, SMC3 and NIPBL ChIP-seq tracks in control and FK866-treated conditions. Blue and red arches at the bottom represent lost or gained loops between treated and control conditions (|log2FC| > 0 & FDR < 0.05) (**A**). Average plots for CTCF (top) and SMC3 (bottom) ChIP-seq experiments on CTCF and SMC3 peaks respectively (**B**). PCA based on EBV RNA-seq reads showing NAMPTi is responsible for most of the changes in gene expression in three biological replicates (**C**). Volcano plot showing all up- (red) and downregulated (blue) genes upon FK866 treatment (|log2FC| > 0 & FDR < 0.05) (**D**). Western Blot showing the decrease in Latent Membrane Protein 1 (LMP1) levels after FK866 treatment (N = 3 biological replicates) (**E**). Barplot showing viral copy number expressed as EBV copies per cell (N = 3 biological replicates) (**F**).

**Figure 7.**
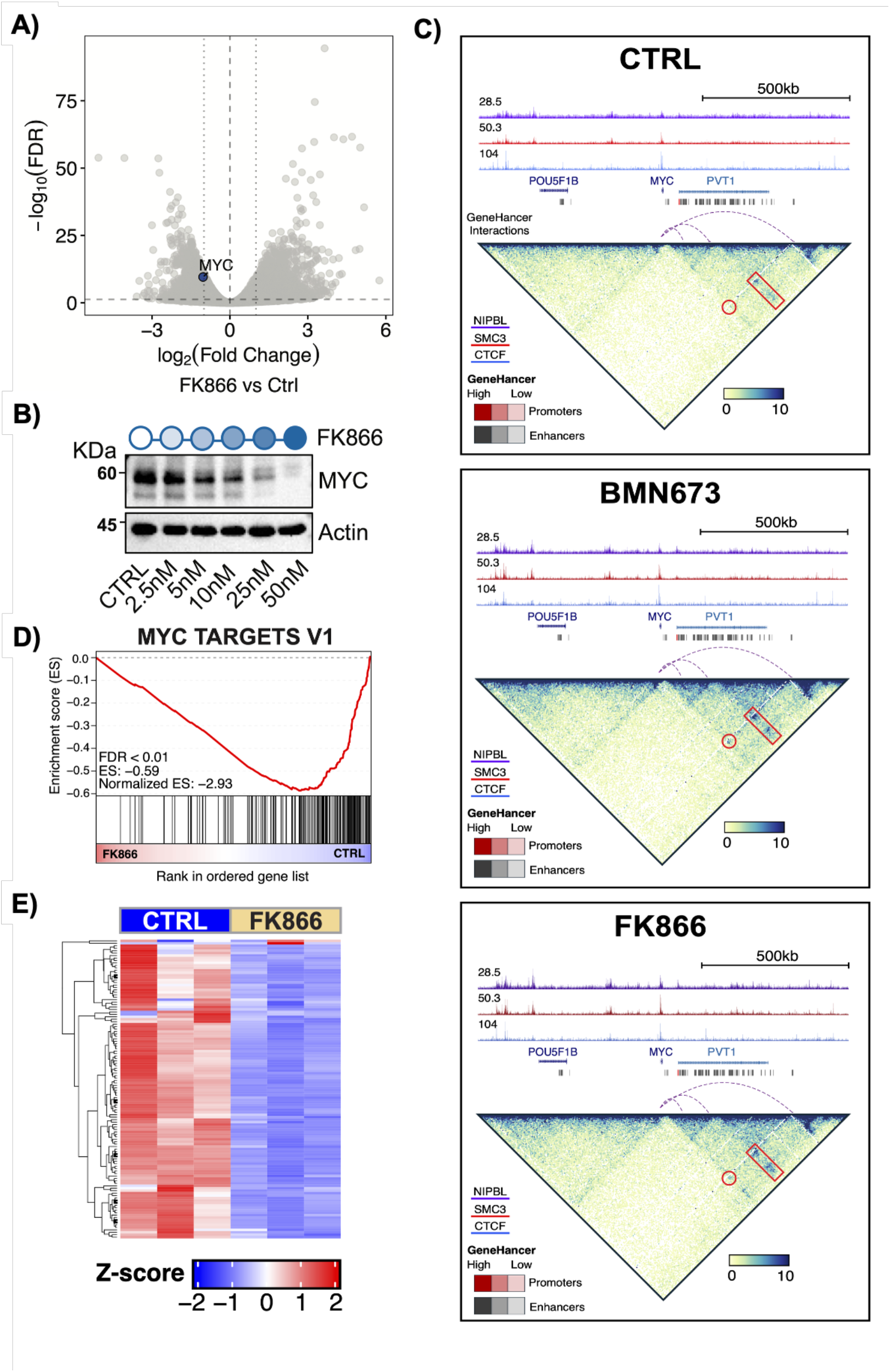
**NAMPTi induces a downregulation of MYC expression through chromatin architecture rearrangement**: Volcano plot showing MYC among the significantly downregulated genes upon FK866 treatment (**A**). Western Blot showing the decrease in MYC protein level at increasing concentration of FK866 (**B**). Contact matrices generated from TopoLink assay of the MYC locus in control, BMN673 and FK866 conditions. On top of each matrix are plotted NIPBL, SMC3 and CTCF ChIP-seq read-depth normalized tracks (top to bottom). GeneHancer predicted interactions between MYC promoter and distal enhancers are also plotted as purple dashed arches. MYC promoter region and all enhancers are indicated as red or gray bars respectively on top of the arches (**C**). GeneSet Enrichment Analysis (GSEA) of RNA-seq revealed MYC pathway to be downregulated (**D**). Heatmap showing the Z-score of genes involved in the GSEA MYC pathway (N = 3 biological replicates) (**E**).

Next, we assessed the effect of inhibition of NAD salvage pathway activity on viral gene expression by analyzing our RNA-seq dataset for EBV transcripts. Principal component analysis (PCA) of the RNA-seq viral transcriptome revealed that FK866 treatment accounted for 53% of the variance in EBV gene expression (**Figure 6C**). Notably, NAD depletion preferentially downregulated latent viral transcripts, including LMP1 and EBNAs (**Figure 6D**). Importantly, *LMP1* is located within a region of the EBV genome that undergoes chromatin structure remodeling following FK866 treatment (**Figure 6A**). Consistent with the RNA-seq data, western blot analysis confirmed reduced LMP1 protein expression (**Figure 6E**). RNA-seq analysis also indicated that FK866 treatment increased expression of a few viral lytic genes (**Figure 6**). To determine whether NAD depletion is sufficient to trigger EBV lytic reactivation, we quantified viral genome copy number in cells before and after FK866 treatment using digital droplet PCR (ddPCR). We observed no significant change in EBV copy number following NAMPT inhibition (**Figure 6F**), indicating that although NAD depletion induces the transcription of lytic viral genes, NAD depletion alone is insufficient in inducing viral replication. These findings suggest that NAD⁺ salvage pathway activity contributes to the regulation of EBV chromatin architecture and viral gene expression.

### NAD Salvage pathway sustains MYC expression in EBV+ B-cells

In LCLs, PARP1 activity has been shown to influence MYC expression^116^. Consistent with this, our RNA-seq analysis revealed a downregulation of MYC transcripts following FK866 treatment (**Figure 7A)**, which was further confirmed at the protein level by western blot analysis when treating with higher doses of FK866 (**Figure 7B**). To determine whether NAMPT inhibition represses MYC expression by altering the 3D chromatin architecture of its locus, we leveraged the TopoLink assay dataset to assess chromatin organization at the MYC locus in LCLs after FK866 treatment. To further evaluate whether NAD depletion phenocopies PARP1 inhibition, we also conducted TopoLink in LCLs treated with the potent PARP inhibitor BMN673^117,118^. We first confirmed by western blot analysis that FK866 treatment inhibits PARylation at levels comparable to BMN673 (**Supplemental Figure 8B**). Next, we established by ChIP-seq that PARP inhibition recapitulates the effects of FK866 treatment on the binding of CTCF, SMC3, and NIPBL in LCLs (**Supplemental Figure 9A).** PARP inhibition altered cohesin occupancy at specific regions similarly to NAD depletion, suggesting a shared mechanism of chromatin dysregulation (**Supplemental Figure 9A).**

TopoLink assay analysis revealed that both FK866 and BMN673 treatments altered the 3D chromatin architecture surrounding the MYC promoter (**Figure 7C**). Both treatments induced a comparable pattern of chromatin loop gains and losses at this locus, with striking predominant frequent interactions with MYC promoter and its distal enhancers, showing a striping pattern in BMN673 and FK866 specific matrices. These changes occurred at regions co-occupied by CTCF and cohesin. While both drugs promoted increased SMC3 binding and destabilized CTCF occupancy, FK866 uniquely enhanced NIPBL binding around the MYC promoter (**Figure 7C, bottom**). Notably, downstream of the MYC promoter, FK866 and BMN673 treatments exhibited opposing effects on chromatin loop formation. This divergence likely reflects differential regulation of CTCF binding, as FK866 treatment led to a loss of CTCF occupancy at one end of the loop, whereas CTCF binding was maintained in BMN673-treated cells (**Figure 7C**). These results suggest that NAD depletion may exert broader effects on chromatin architecture beyond those caused by PARP inhibition alone, potentially through NIPBL stability or function. In addition, our data indicated that MYC expression is responsive to changes in NAD levels in B-cells. Since MYC is an important EBV target and frequently deregulated in B-cell malignancies, we next determined whether reduced MYC activity impacts the expression of its target gene programs after NAD depletion. Gene Set Enrichment Analysis (GSEA) of RNA-seq data revealed that MYC target genes were significantly enriched among those downregulated following FK866 treatment (**Figure 7D**) and FK866 led to robust repression of MYC target gene expression (**Figure 7E**), indicating that alterations in chromatin looping at the MYC promoter contribute directly to the transcriptional response triggered by NAD depletion.

To further examine the relationship between altered chromatin protein occupancy and the MYC transcriptional signature, we compared the MYC Targets V1 and V2 GSEA gene sets with genes annotated within 3 kb of ChIP-seq peaks assigned to each cluster (**Supplemental Figures 9A** and **9B**). MYC Targets V1 genes showed greater proportional representation in cluster 1, accounting for 100 of 7,849 associated genes (1.27%), compared with 40 of 7,628 in cluster 2 (0.52%) and 115 of 17,560 in cluster 3 (0.65%). Thus, cluster 1 contained nearly as many MYC target genes as cluster 3 despite being associated with fewer than half as many genes overall. This distribution is consistent with a preferential association of MYC target genes with cluster 1 sites. Overall, these results demonstrate that NAMPT inhibition is associated with altered chromatin architecture at the MYC locus, reduced MYC expression, and repression of MYC-associated transcriptional programs. Overall, our findings support a role for NAD⁺ salvage pathway activity in maintaining chromatin organization and MYC-associated oncogenic transcriptional programs in EBV-transformed B cells.

### NMN Supplementation attenuates Chromatin Architectural Changes Induced by NAMPT Inhibition

To further investigate the effects of NAD⁺ salvage pathway inhibition on chromatin organization, we performed H3K27ac HiChIP in LCLs treated with FK866 alone, or in combination with nicotinamide mononucleotide (NMN) supplementation. NMN supplementation was included to bypass NAMPT inhibition and replenish NAD⁺, allowing us to assess whether the effects of FK866 on chromatin contacts could be reversed by restoring NAD⁺ availability.

We analyzed 1,502 loops that differed from control in both treatment conditions (P < 0.05 in each comparison) and changed in the same direction. Within this subset, NMN supplementation reduced the magnitude of the changes, bringing loop intensities closer to control values. The paired difference in absolute log_2_ fold change, calculated as |log_2_FC(FK866)| − |log_2_FC(FK866+NMN)|, was shifted toward positive values (median = 0.047; two-sided Wilcoxon signed-rank P < 2×10^-16^; **Figure 8A, left**). The mean absolute log_2_ fold change decreased from 0.885 (95% CI, 0.868–0.902) with FK866 alone to 0.822 (95% CI, 0.806–0.839) with FK866 plus NMN (**Figure 8A, right**). These results demonstrate partial attenuation of FK866-associated contact changes by NMN supplementation within this set of interactions. In addition, 76.3% of the anchors associated with these loops overlapped an SMC3 ChIP-seq peak (**Figure 8B**), indicating that the contacts frequently involved cohesin-occupied regions.

**Figure 8.**
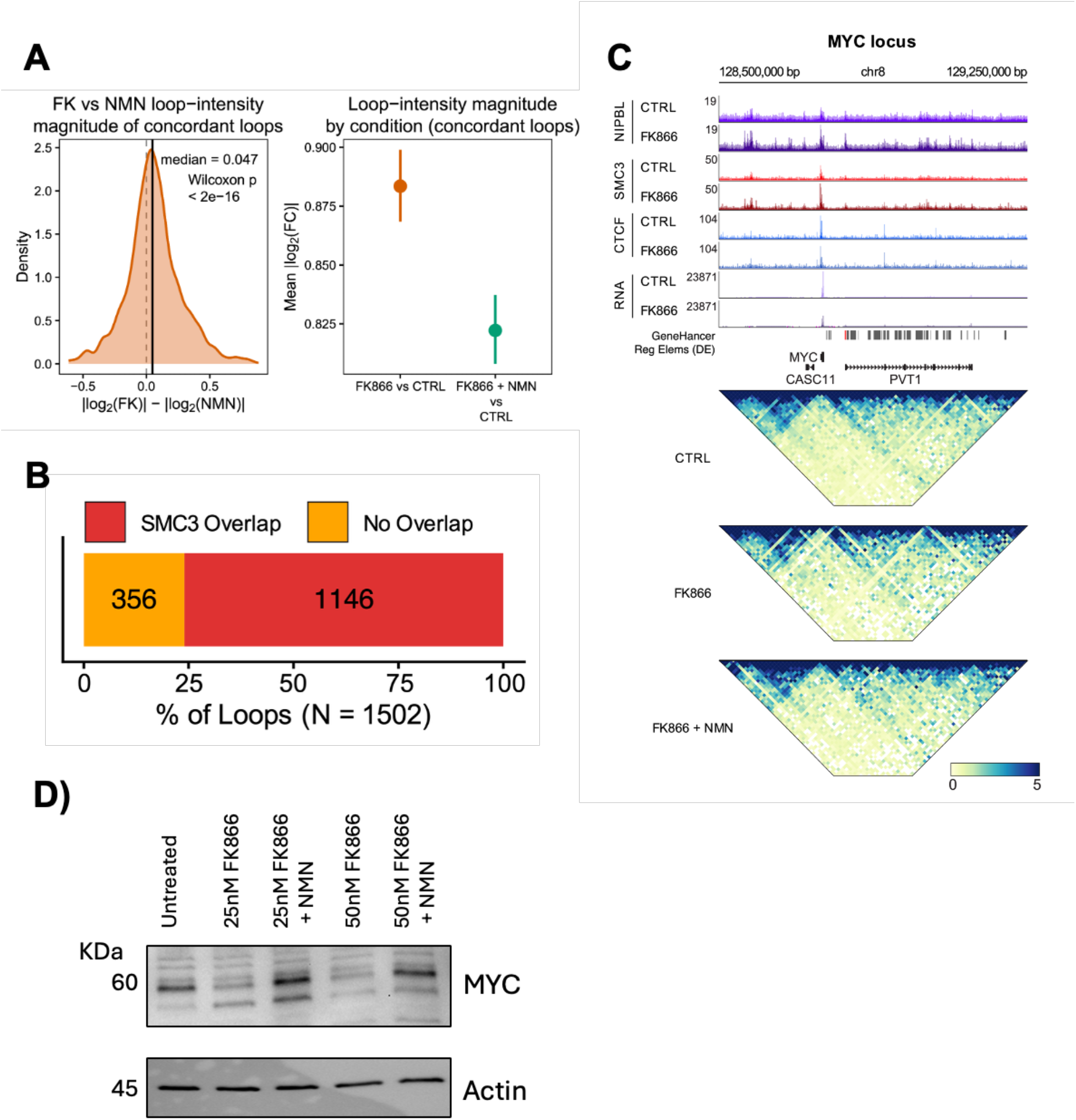
**NMN supplementation partially restores chromatin interactions**: Loop-intensity changes in concordant differential loops, defined as loops called differential in both FK866 vs. CTRL and FK866+NMN vs. CTRL and changing in the same direction (n = 1502). Density of the per-loop difference in effect magnitude, |log2FC(FK866)| - |log2FC(NMN)|, positive values indicate a smaller change upon NMN co-treatment. Dashed line, median (0.047), solid line, zero. P-value was calculated using a two-sided Wilcoxon signed-rank test against a null median of zero (**A**, left). Mean absolute loop-intensity change per comparison (points, mean |log2FC|; error bars, [95% confidence interval/SEM]) (**A**, right). Fraction of concordant differential loops with at least one anchor overlapping an SMC3 ChIP-seq peak (n = 1502 loops) (**B**). MYC locus, from top: NIPBL, SMC3 and CTCF ChIP-seq and RNA-seq coverage in control and FK866-treated cells; group-autoscaled track maxima are indicated at left. GeneHancer regulatory elements are shown below, followed by the RefSeq genes MYC, CASC11 and PVT1. Bottom, H3K27ac HiChIP contact maps for control, FK866 and FK866+NMN over the same interval, 10-kb resolution, read-depth- and KR-normalized; color scale, [normalized contact frequency], 0–5 (**C**). Western blot showing recovery of MYC protein levels after daily supplementation with 100uM NMN after high dose FK866 treatment (**D**).

**Figure 9.**
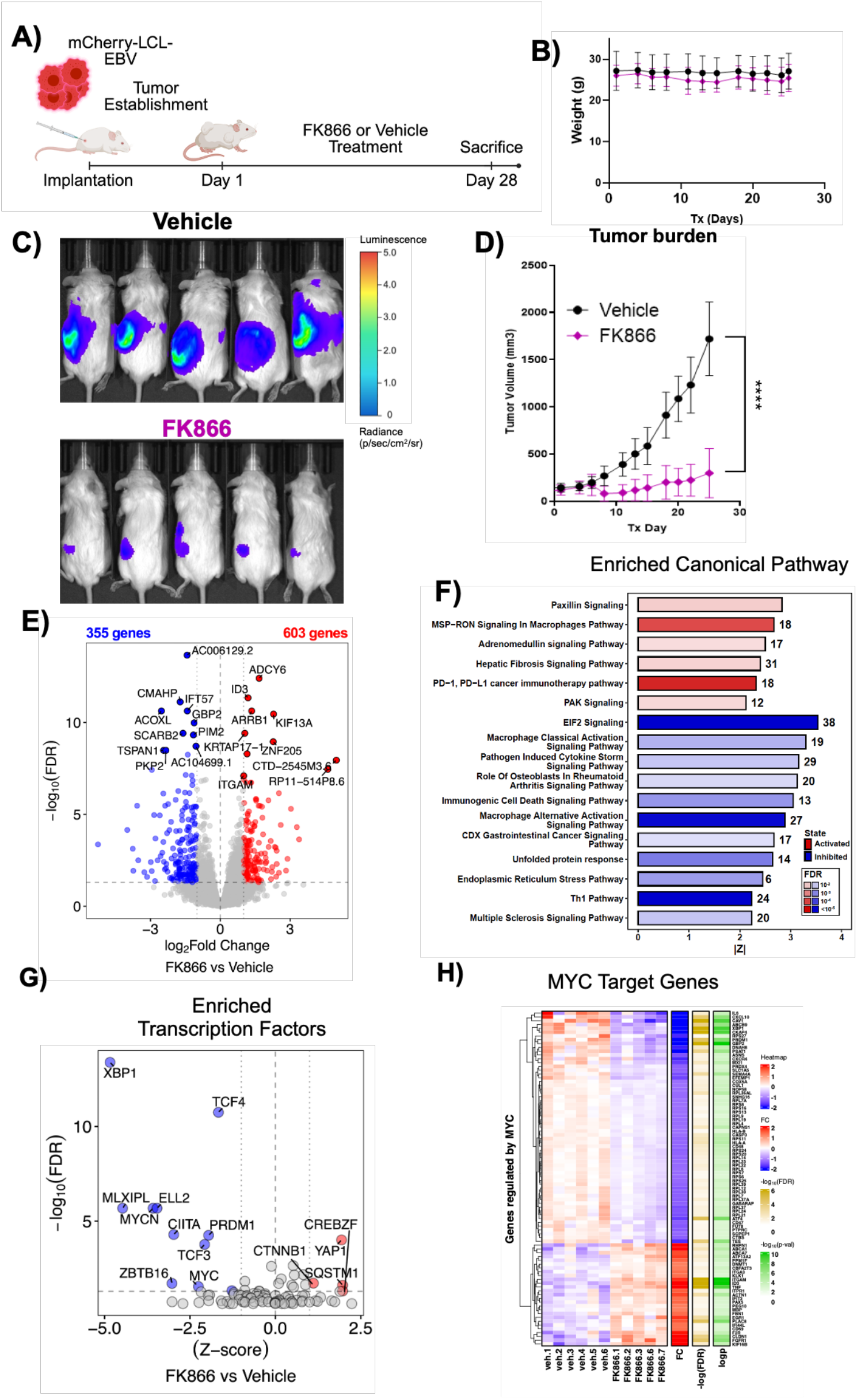
**In vivo treatment of EBV-positive tumors with FK866 is effective in reducing tumor size**: Cartoon describing the treatment procedure of the in vivo study. Mice (five males and five females per treatment group) were subcutaneously engrafted with \(1 \times 10^6\) NHC1 LCLs expressing mCherry and enhanced luciferase (eLUC). On day 5 post-engraftment, mice were randomized into treatment groups based on baseline bioluminescent signal. Tumor burden was monitored longitudinally using the IVIS imaging system following intraperitoneal injection of D-luciferin (7.5 mg/kg), with total-body bioluminescence quantified as photon flux (photons/sec) (**A**). Lineplot showing the weight of the mice during treatment as a proxy of animals well-being (**B**). (**C**). Lineplot displaying the tumor size over time (**** = P-value < 0.0001) (**D**). Volcano plot showing the differentially expressed genes in the tumors after FK866 treatment. In blue are indicated the significantly downregulated genes, in red the upregulated genes while in grey the nonsignificant ones (|log_2_FC| > 1 & FDR < 0.05) (**E**). Barplot showing the activated (red) or inhibited (blue) enriched canonical pathways identified by IPA (**F**). Volcano plot displaying the inhibited (blue) or activated (red) transcription regulators as determined by IPA (**G**). Heatmap showing the Z-score, log_2_FC, −log_10_(FDR) and –log_10_(P-value) of genes that constitute the GSEA MYC TARGETS V1 pathway (**H**).

We next examined the MYC locus, given the MYC target signatures identified above and the established role of this oncogene in EBV-driven B-cell proliferation. Consistent with transcriptional and HiC data, H3K27ac HiChIP revealed altered contacts between the MYC promoter and distal enhancers following FK866 treatment, which were attenuated by NMN supplementation (**Figure 8C**). In addition, NMN supplementation following high-dose FK866 treatment also restored MYC protein levels (**Figure 8D**). Together, these findings support a link between NAD⁺-dependent maintenance of chromatin interactions across the MYC locus and MYC protein expression in EBV-transformed B cells.

### Inhibition of NAD salvage pathway activity hampers EBV-driven tumor growth in vivo

MYC plays a critical role in driving lymphomagenesis. Our recent work showed that PARP inhibition downregulated MYC expression in vivo, inhibiting the grow of EBV-driven malignancies^116^. Our data showing that the NAMPT inhibitor FK866 epigenetically represses MYC expression suggests that NAD depletion may also impair EBV-driven tumor growth. To test this hypothesis, 16 NSG mice (8 females and 8 males) were engrafted with luciferase-expressing EBV+ lymphoblastoid cells to enable in vivo monitoring of tumor progression via bioluminescence imaging (**Figure 9A**). Seven days post-engraftment (T0), mice were stratified based on average tumor flux (photons/second [p/s]) and randomized into two groups (n=8 per group, 4 females and 4 males) to receive either FK866 or vehicle (Veh) by daily oral administration. Mice were monitored daily, and no significant differences in body weight or overall health were observed between groups during treatment (**Figure 9B**). At day 28 (T28), mice were sacrificed and tumors collected. Bioluminescence analysis revealed a significant reduction in total tumor flux in the FK866-treated group compared to controls (**Figures 9C** and **9D**). These results demonstrate that NAD salvage pathway inhibition restrains EBV-driven tumor growth in vivo. To determine whether the in vivo NAMPT effects are due to repression of oncogenic programs such as MYC, we performed RNA-seq analysis on tumors from FK866- and vehicle-treated mice.

Comparative transcriptomic analysis revealed 958 differentially expressed genes in FK866-treated tumors, including 603 upregulated and 355 downregulated transcripts (**Figure 9E**). Pathway enrichment analysis identified activation of Paxillin and PAK signaling cascades — associated with cytoskeletal remodeling and immunological synapse formation — alongside engagement of the PD-1/PD-L1 immune checkpoint pathway and suppression of key proliferative (E2F), pro-inflammatory, unfolded protein response (UPR), and immunogenic cell death (ICD) programs (**Figure 9F**). This composite signature suggests that FK866 treatment induces proliferative arrest and cytoskeletal reorganization, contributing to reduced tumor growth. Interesting, pathway analysis further revealed a significant inhibition of cellular signaling pathway connected to multiple sclerosis (**Figure 9F**), which has been recently associated with EBV infection^119^, suggesting a potential inhibitory role of NAD depletion on EBV-associated autoimmunity.

Importantly, FK866-treated tumors exhibited a transcriptional signature that was indicative of activation of a few regulators associated with distinct but complementary cellular functions in B-cells, including YAP1 and CTNNB1 (β-catenin), key effectors of the Hippo and Wnt pathways, respectively, which are linked to stress adaptation and cell survival (**Figure 9G**). YAP1 activation is consistent with its recently established role in EBV infection, where YAP1 cooperates with viral proteins such as LMP1 to regulate EBV infection in epithelial cells^120–123^. On the other hand, the effect of NAD depletion led to the inhibition of multiple key transcriptional regulators including TCF3, MYC, MYCN, CIITA, ZBTB16, ELL2, MLXIPL, and XBP1 (**Figure 9G**). This pattern of repression is consistent with a global shutdown of B-cell identity and function. Consistent with our in vitro findings, MYC target genes were significantly downregulated in FK866-treated tumors. (**Figure 9H**), supporting the conclusion that NAD salvage pathway activity is essential for maintaining MYC function and MYC-driven tumor growth in B-cells. Similarly, expression of XBP1 target genes, crucial for ER stress adaptation and survival and plasma cell differentiation^124–127^, was significantly reduced in FK866-treated tumors (**Supplemental Figure 10**), indicating that the effect of NAD depletion may target multiple transcriptional pathways that are essential for tumor growth. Collectively, these data support a model in which inhibition of NAD salvage pathway activity disrupts CTCF/cohesin looping, leading to impaired regulation of MYC and attenuation of MYC-driven transcriptional programs (**Figure 10**).

**Figure 10.**
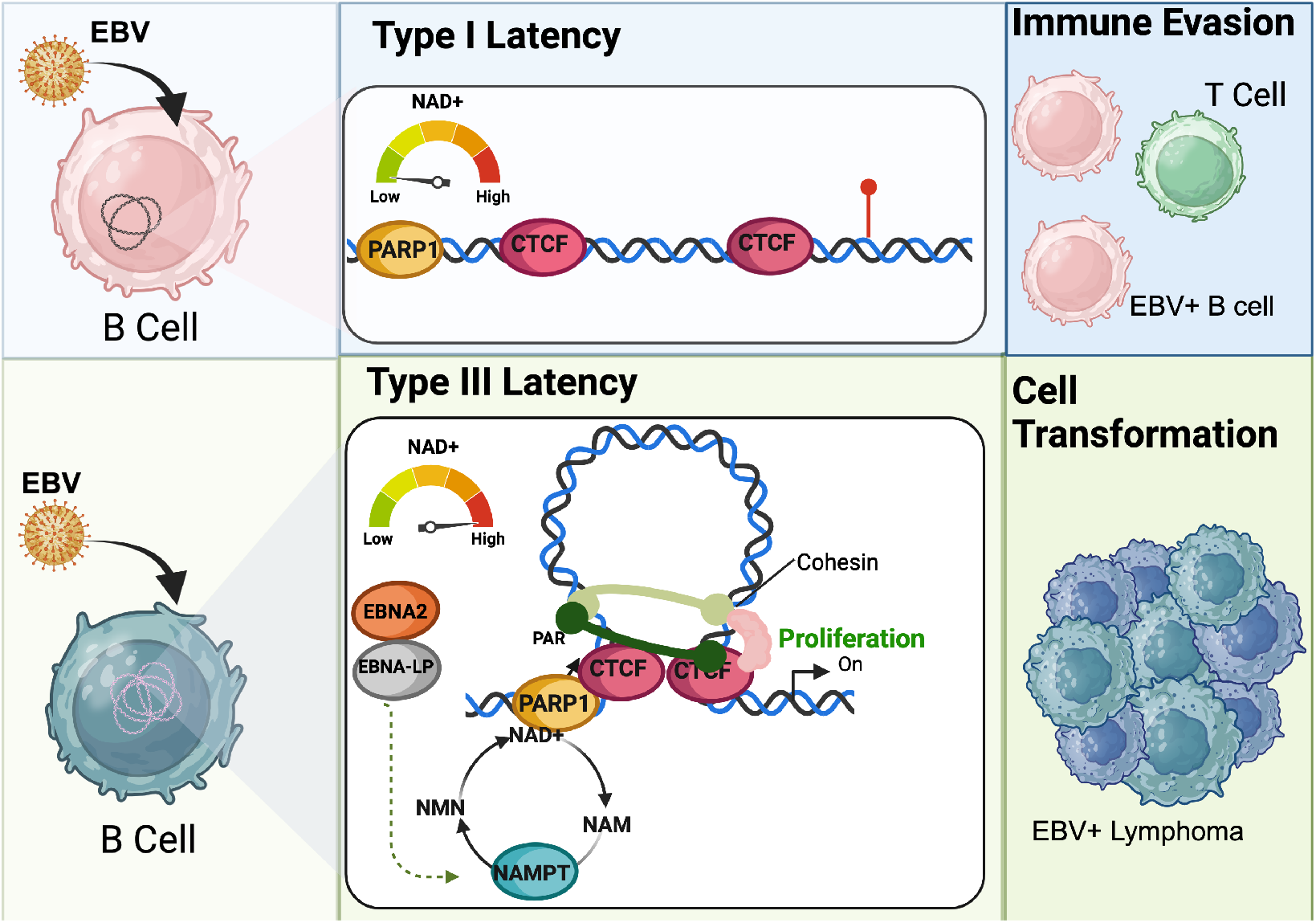
**EBV latency III is responsible for the activation of the NAD salvage pathway necessary for latency maintenance and B-cell transformation**: Schematic model showing how EBV type III latency is responsible for the activation of the NAD salvage pathway through EBNA2 and EBNA-LP which, in turn, is necessary to sustain higher PARP1 activity that helps shaping chromatin architecture to achieve B-cell transformation.

## Discussion

Our study identifies NAD⁺ salvage as a metabolic dependency in EBV-driven B-cell transformation and tumor growth and links this pathway to host chromatin regulation. EBV infection of primary B cells increased expression of NAMPT, the rate-limiting enzyme in NAD⁺ salvage, while NAMPT inhibition with FK866 drug impaired B-cell immortalization and caused pronounced cytotoxicity in LCLs at higher doses. These findings support the metabolic rewiring toward reliance on NAD⁺ salvage during infection and in transformed cells. Our analysis indicates that NAMPT induction coincided with peak expression of latent viral proteins, suggesting that the viral latency program contributes to this response. Consistent with this possibility, EBNA2 and EBNA-LP occupied the NAMPT promoter and a distal regulatory region characterized by H3K27ac enrichment and chromatin contacts with the promoter, suggestive of enhancer activity. Indeed, CRISPR interference targeting this region reduced NAMPT protein levels, establishing the contribution EBNA2/EBNA-LP occupied region to NAMPT expression. Given the established roles of EBNA2 and EBNA-LP in regulating host transcription and chromatin, these observations support a model in which EBV promotes NAD⁺ salvage through regulation of the NAMPT locus. The specific contributions of these viral proteins to NAMPT induction however remain to be established.

Recent work by Müller-Durovic and colleagues demonstrated that EBV infection induces NAD biosynthesis via the de novo pathway and IDO1 activation during the early stages of infection^40^. While our findings are consistent with this initial increase in NAD levels, we extend these observations by revealing a critical role for the NAD salvage pathway, through NAMPT upregulation, in sustaining NAD production and supporting EBV-driven cellular reprogramming. Overall, our findings contribute to position NAD metabolism as a key axis in the molecular interplay between EBV and its host. Mechanistically, we show that NAMPT-dependent NAD^+^ salvage pathway activity is critical during the early phase of EBV infection and remains required in established latent infection for the maintenance of chromatin and transcriptional programs associated with EBV-transformed B cells. Pharmacological inhibition of NAMPT with FK866 reduces intracellular NAD^+^ abundance and induces widespread transcriptional and chromatin changes. These effects are consistent with reduced activity of NAD^+^-dependent enzymatic processes, including PARP1 activity. We previously demonstrated that EBV infection activates PARP1, and that PARP1 activity is essential for establishing a permissive chromatin landscape across the EBV episome, thereby facilitating and sustaining viral gene expression^18–20^. Because PARP1 requires NAD^+^ as a co-substrate, increased activity of the NAD^+^ salvage pathway may help maintain the NAD^+^ supply needed at least for PARP1-dependent regulation of viral chromatin.

The importance of NAD⁺ availability may extend beyond the regulation of viral chromatin. Previous work has demonstrated that EBV-transformed cells are highly dependent on PARP1 for DNA repair^105^, highlighting a potential vulnerability arising from the metabolic demands of maintaining EBV latency. Consistent with this possibility, our transcriptomic analyses revealed deregulation of DNA repair-associated pathways following NAMPT inhibition. Thus, reduced PARP1 activity may account for some of the observed transcriptional changes, while others could reflect broader effects of NAD⁺ depletion on chromatin regulation and DNA repair in EBV-transformed cells.

Our findings suggest that NAD⁺ availability contributes to chromatin organization by influencing the distribution and function of architectural proteins. NAMPT inhibition was accompanied by compartment switching, sharper domain boundaries, reduced contacts between distant domains, and a tendency toward reinforcement of pre-existing contacts. Increased SMC3 and NIPBL occupancy within this largely preserved framework raises the possibility that NAD⁺ depletion shifts the balance between cohesin recruitment, retention, and redistribution. The increased local retention of cohesin complex we observed could favor intradomain contacts while limiting or redirecting longer-range interactions, potentially uncoupling cohesin accumulation from its architectural activity. A precedent comes from NIPBL-depletion studies, in which cohesin-STAG1 accumulates at CTCF sites despite impaired genome folding. The largely unchanged total abundance of CTCF and SMC3 further supports regulation through chromatin association or function, although the effects on cohesin residence time, translocation, and extrusion ability require further direct testing.

The transcriptional consequences of this response may depend on genomic context. CTCF motif enrichment in accessible regions in control cells and predicted CTCF regulatory activity among FK866-responsive cluster 1 genes capture distinct regulatory features. At cluster 1 sites, the transcriptional signature accompanied increased SMC3 and NIPBL occupancy with relatively low and largely stable CTCF binding, suggesting that changes in cohesin association could influence transcription without increased CTCF occupancy. Such changes may selectively affect regulatory interactions, as illustrated by the accompanying alterations in MYC promoter–enhancer contacts and protein expression. NMN-mediated attenuation of contact changes and restoration of MYC protein levels provide complementary support for a relationship between NAD⁺ availability, local chromatin organization, and transcriptional output, while demonstrating that these effects are at least partially reversible.

Our results also indicate that NAD⁺ availability and local chromatin organization play an important role in the transcriptional output of the MYC locus, where altered promoter–enhancer contacts were accompanied by reduced MYC transcription and suppression of its downstream gene expression program following NAMPT inhibition. Given the central role of MYC in EBV-driven B-cell proliferation128–130, these findings provide a potential link between local chromatin changes and reduced growth. These observations suggest that NAD⁺ availability contributes to the chromatin environment supporting MYC expression. They also illustrate how changes within an existing architectural framework may influence oncogenic transcription without requiring widespread formation or loss of chromatin loops. MYC suppression occurred alongside broader stress, checkpoint, and antiviral transcriptional signatures, consistent with a shift away from EBV-sustained proliferation. These responses may reflect direct effects of NAD⁺ depletion or secondary consequences of altered chromatin organization and genome maintenance, consistent with the enrichment of genes involved in DNA strand elongation and mismatch repair among those differentially expressed following NAMPT inhibition.

As predicted by our studies in cell lines, targeting NAMPT and NAD salvage pathway hamper EBV⁺ lymphoma growth in xenografts model of EBV malignancies in vivo. Reduced tumor burden was accompanied by repression of MYC and XBP1 target programs and checkpoint-associated transcriptional responses, broadly recapitulating the changes observed in cultured cells. The consistent suppression of MYC-associated transcription across these settings suggests that weakening this program may contribute to tumor control. Similar responses to PARP inhibition support a contribution from NAD⁺-dependent chromatin regulation, alongside other metabolic consequences of NAMPT inhibition. These findings identify NAD⁺ salvage as a vulnerability in EBV-associated lymphoma and provide a rationale for investigating whether the relationship between NAD⁺ availability, MYC regulation, and tumor growth extends to other MYC-driven malignancies.

The effects of NAD⁺ depletion extended to the EBV episome, where altered chromatin contacts and increased SMC3 occupancy accompanied repression of LMP1 and EBNA transcripts. These findings suggest that NAD⁺ salvage supports the viral chromatin environment associated with growth-promoting latent gene expression. Unlike the predominant host response, CTCF occupancy decreased across the viral genome, raising the possibility that viral factors or the episomal chromatin context confer distinct requirements for CTCF recruitment or retention. Changes involving oriLyt and the BZLF1 promoter also connect NAD⁺ availability to regions implicated in the latent–lytic switch. Although MYC regulates these interactions and can suppress EBV reactivation, MYC repression and altered viral contacts were not accompanied by increased BZLF1 expression under our sublethal treatment conditions. These observations raise the possibility that NAD⁺ depletion weakens the latent transcriptional program and shifts viral chromatin toward a state more permissive to reactivation, although full priming of viral chromatin for lytic reactivation may require more pronounced NAD⁺ inhibition and MYC depletion than that achieved under the sublethal treatment conditions used here. In addition, changes in mismatch-repair-associated transcription could have additional implications for the viral life cycle, as host mismatch repair factors are recruited to EBV replication compartments and have been implicated in the maturation of newly synthesized viral genomes. NAD⁺ depletion may therefore influence both the chromatin regulation of viral transcription and the cellular repair machinery available during productive replication. Its effects on mismatch repair activity, viral genome maturation, and progression through the lytic cycle require direct testing.

Some limitations should be considered when interpreting our findings. Our Hi-C analyses identified significant changes in larger-scale chromatin interactions that were consistent with the cohesin occupancy and transcriptional findings, although the resolution of these data limited assessment of smaller loops. Our functional studies also relied primarily on pharmacological NAMPT inhibition with FK866. The sublethal concentration allowed molecular effects to be examined without detectable changes in LCL viability, proliferation, or cell-cycle progression. NMN rescue and complementary metabolomic findings with a NAMPT-targeting PROTAC support the involvement of NAMPT-dependent NAD⁺ salvage, while genetic NAMPT perturbation would provide independent validation of its contribution to the chromatin and transcriptional phenotypes. The extent of reversibility requires further studies, as NMN supplementation attenuated, but did not fully reverse, the HiChIP-detected contact changes, potentially reflecting differences in the kinetics of chromatin responses to NAMPT inhibition and NAD⁺ replenishment. In addition, although our interpretation focused on PARP1, NAD⁺ is a co-substrate for other chromatin regulators, including sirtuin deacetylases131–133, whose contributions cannot be distinguished by our proteomic and chromatin-accessibility analyses. Finally, comprehensive metabolic and chromatin profiling was performed mainly in established EBV-transformed cells. Although NAMPT inhibition impaired EBV-driven B-cell immortalization, the metabolic requirements of initial infection and transformation may differ from those sustaining established latency.

Our findings identify NAMPT-dependent NAD⁺ salvage as a metabolic dependency in EBV-transformed B cells and link NAD⁺ availability to cohesin-associated chromatin organization and MYC expression. These results suggest that metabolic reprogramming supports the host and viral transcriptional programs sustaining the transformed state. The inhibition of tumor growth following NAMPT blockade supports further investigation of NAD⁺ salvage as a therapeutic vulnerability in EBV-associated malignancies.

## Acknowledgements

Laura Corveleyn acknowledges the Research Foundation Flanders – FWO for personal funding (1SF2622N) and awarding a mobility grant to join the Sidoli Lab (V400623N). IT, PL and BG were supported by P01CA269043. IT was also supported by R01AI130209 and P30CA010815. BEG was also supported by R01DE033907. The Sidoli lab gratefully acknowledges for funding the Hevolution Foundation (AFAR), the Einstein-Mount Sinai Diabetes center, and the NIH Office of the Director (S10OD030286). Funding support for The Wistar Institute Proteomics and Metabolomics Shared Resource was provided by Cancer Center Support Grant P30CA010815, and the acquisition of the Thermo Scientific Q Exactive HF-X mass spectrometer was supported by S10 OD023586. H-Y.T. is supported by R50 CA221838. R.S. was supported by T32CA009171, and T32CA288354.

Figure were Created in BioRender.

## Author contributions

Conceptualization: D.M., I. T., R. S.; Data curation: D.M., R.S., S.P.A., P.P., S.S., A.K., A.G., H.T.; Formal analysis: D.M., R.S., Sa.S., S.S., A.K.; Funding acquisition: I.T., B.G., P.L.; Investigation: D.M., R.S., L.B.C., Sa.S., G.N., L.C., A.G., H.T., Y.L.; Project administration: I.T.; Supervision: I.T.; Validation: D.M., R.S., L.B.C.; Visualization: D.M., R.S., I.T.; Writing – original draft: D.M., I.T.; Writing – review & editing: D.M., R.S., I.T., Sa.S., S.S., A.G., B.G., P.L.

## Declaration of interests

The authors declare no competing interests.

---

**Supplemental Figure 1.**
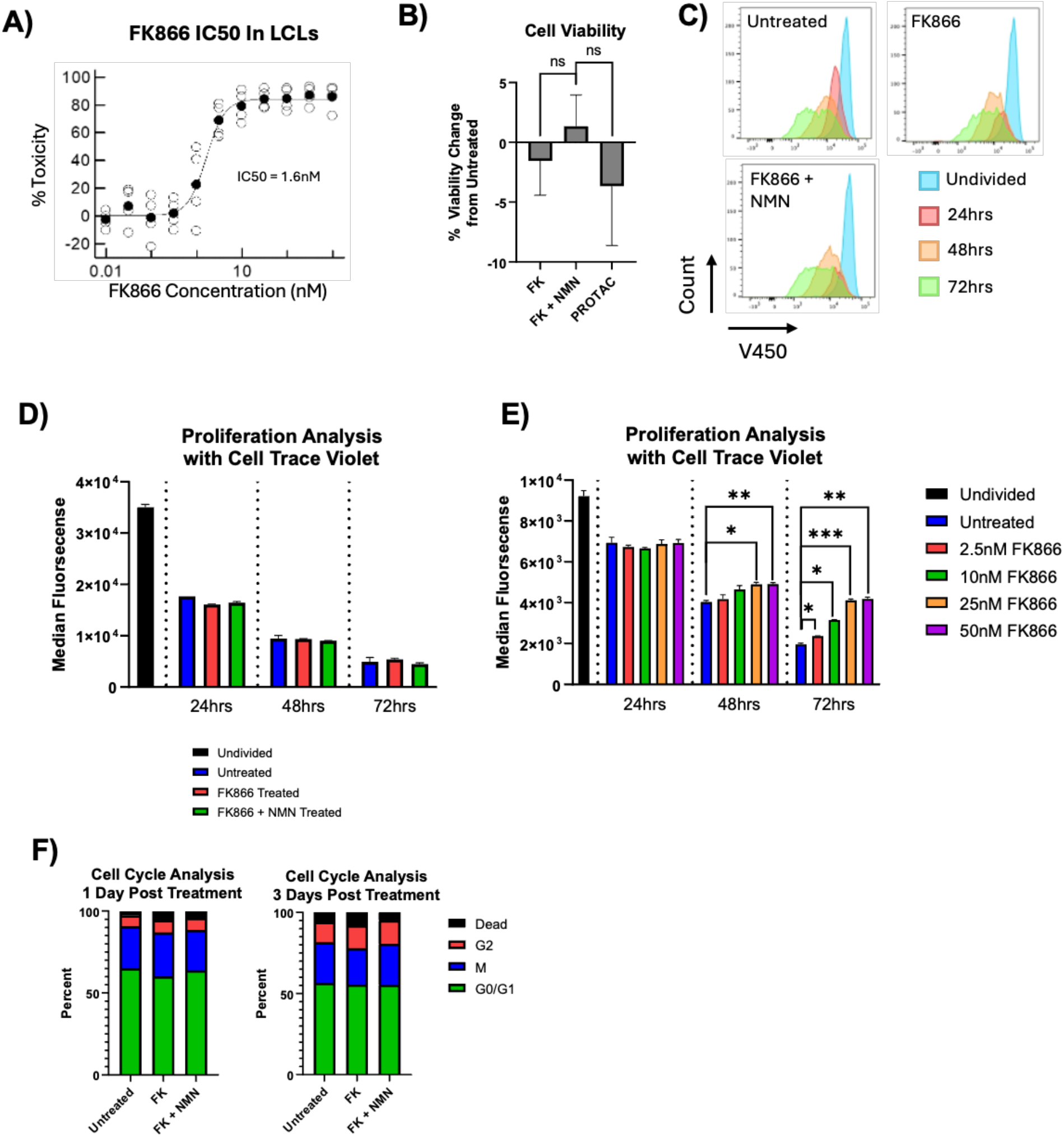
**Sublethal Treatment with FK866 No Effect on Cell Viability or Cell Proliferation**: Determination of FK866 IC₅₀ in LCLs. LCLs were treated with increasing concentrations of FK866 for 72 hours, after which cell viability was quantified using the CellTiter-Glo Luminescent Cell Viability Assay. Luminescence values, which are proportional to cellular ATP content and metabolically active cell number, were normalized to vehicle-treated controls and plotted as percent viability. (**A**). Sublethal FK866 treatment does not significantly reduce cell viability in LCLs. LCLs were treated with 2.5nM FK866, 2.5nM FK866 + 100uM NMN, or 50nM of a NAMPT-targeting PROTAC for 3 days, stained with Zombie NIR Fixable Viability Dye, and analyzed by flow cytometry. Viability was quantified as the percentage of Zombie NIR-negative (live) cells and change in viability from untreated cells is shown. Statistical significance was determined using a two-tailed unpaired t-test with Welch’s correction. (**B**). Cell proliferation assessed by CellTrace Violet dilution. Cells were labeled with CellTrace Violet prior to treatment, fixed at 0, 24, 48, and 72 hours, and analyzed by flow cytometry. Representative overlaid histograms of CellTrace Violet fluorescence are shown (**C**). Proliferation from C was quantified as the median CellTrace Violet fluorescence intensity at each time point. (**D**). Higher doses of FK866 were used to also access cell proliferation using CellTrace Violet. Statistical significance was determined using a two-tailed unpaired t-test with Welch’s correction (**E**).Cell cycle analysis by propidium iodide staining. Cells were fixed, stained with propidium iodide (PI), and analyzed by flow cytometry to determine DNA content. The percentage of cells in the G₀/G₁, S, and G₂/M phases of the cell cycle was quantified and is presented as mean ± SEM from n = X independent experiments.(**F**).

**Supplemental Figure 2.**
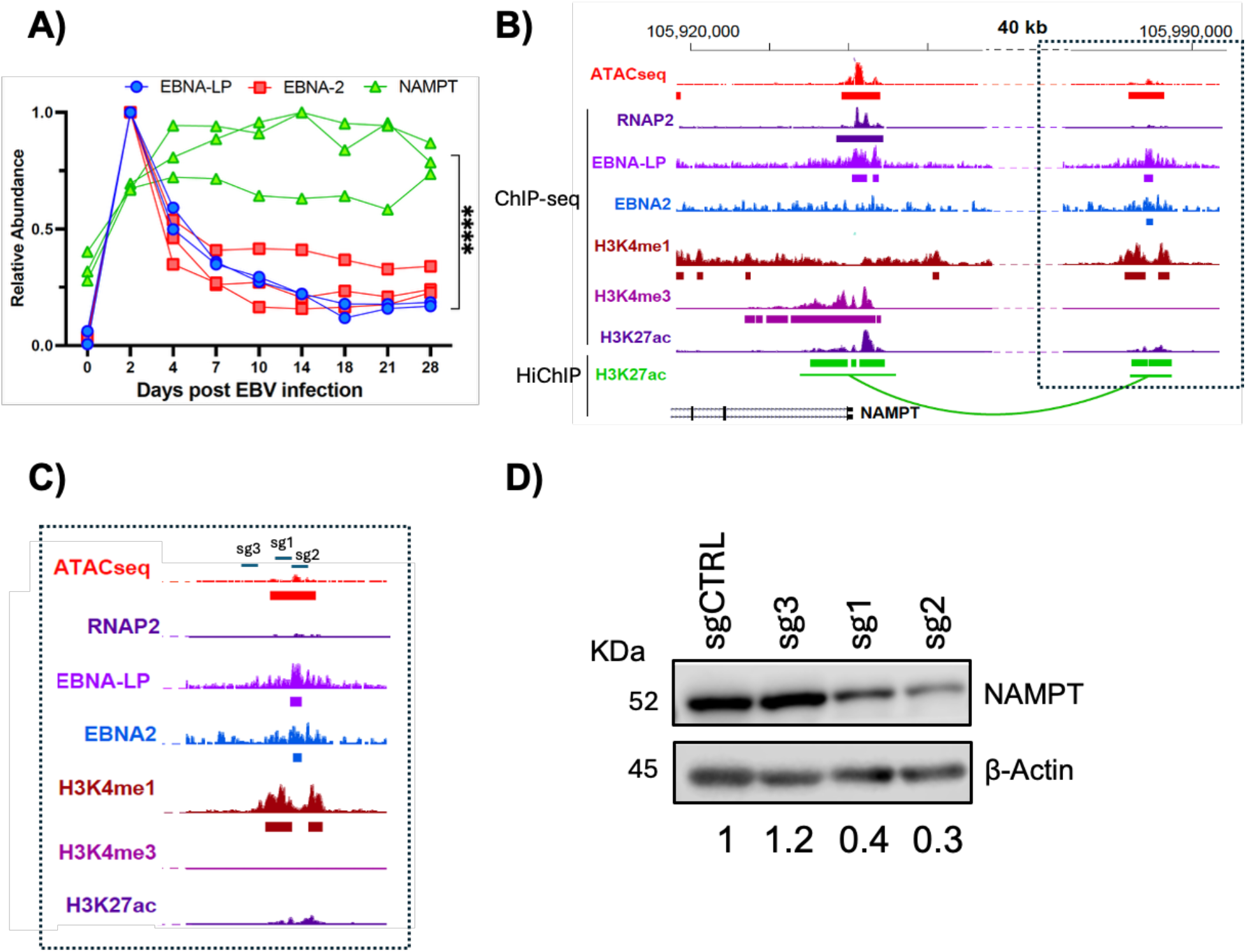
**EBNA2 and EBNA-LP bind regulatory regions at the NAMPT locus**: NAMPT, EBNA2 and EBNA-LP protein abundance during the time course of B-cell infection (**A**). UCSC Genome Browser session showing ATACseq, RNAP2, EBNA-LP, EBNA2, H3K4me1, H3K4me3 and H3K27ac ChIP-seq tracks and H3K27ac HiChIP interactions for NAMPT promoter and distal enhancer **(B).** UCSC Genome Browser session showing binding sites of guide RNAs **(C).** Western blot showing that KO of the EBNA2 and EBNA-lp binding regions of the distal NAMPT enhancer reduces NAMPT expression. sgCTRL is a scramble control, sg3 is a control outside of the binding region, and sg2 and sg2 are within the binding region. ImageStudio quantification software used to quantify relative protein levels **(D).**

**Supplemental Figure 3.**
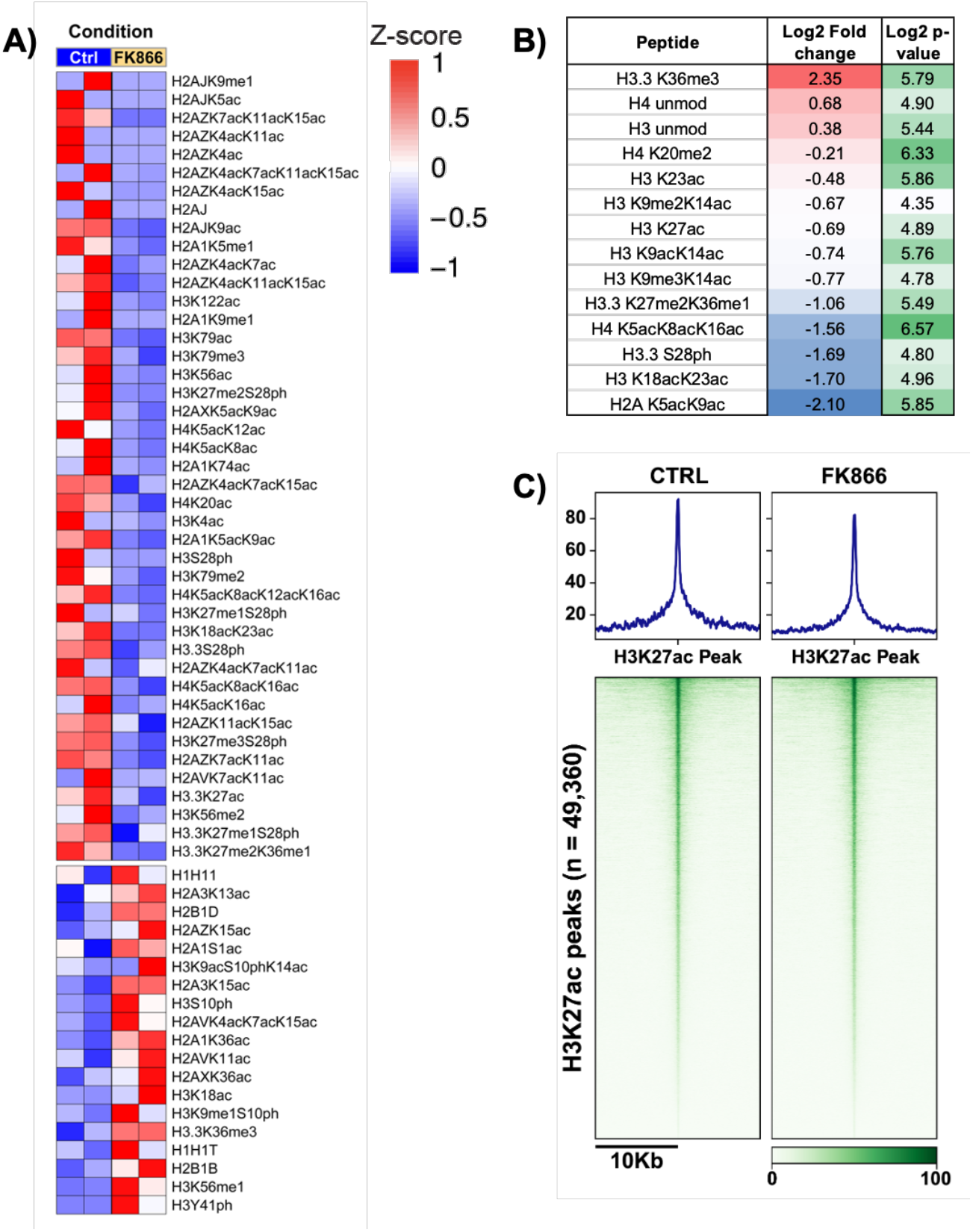
**NAMPTi induces global changes in histone modifications**: Heatmap showing Z-score for each histone modification detected in our LS-MS experiment (**A**). Table showing the top up- (red) and downregulated (blue) histone modifications (**B**). Heatmaps displaying CUT&TAG signal for H3K27ac histone mark genome-wide before and after FK866 treatment (**C**).

**Supplemental Figure 4.**
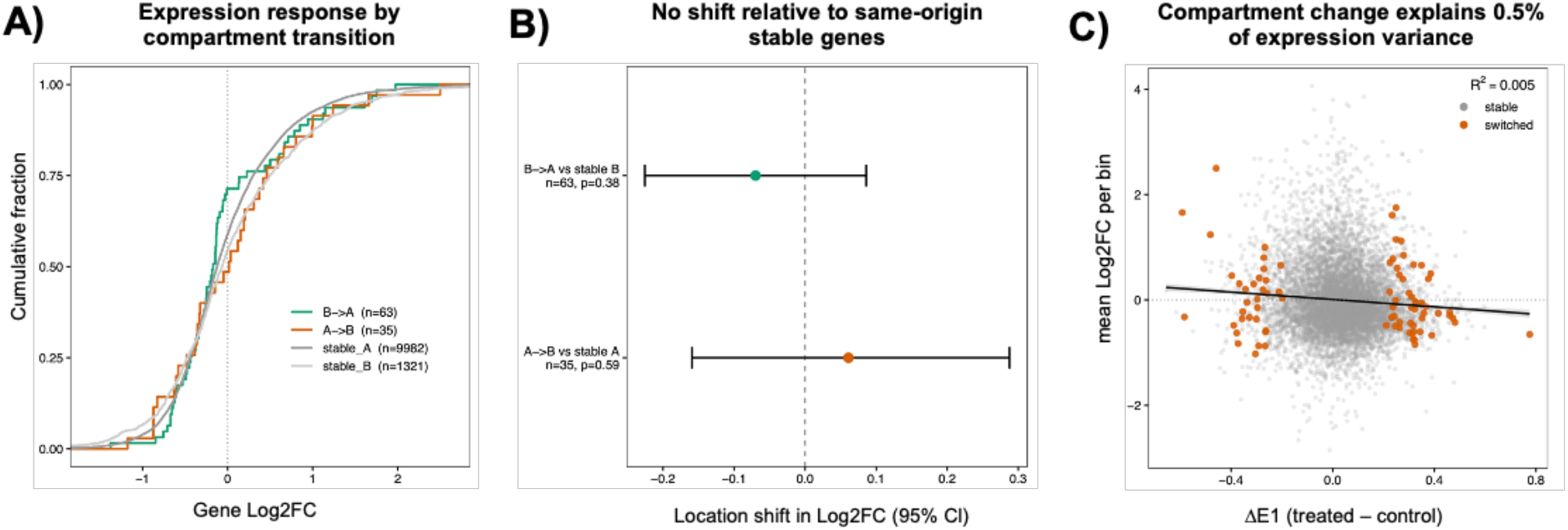
**Compartment switching under NAD salvage inhibition is not associated with coordinated changes in gene expression**. Empirical cumulative distribution functions of gene-level log2 fold changes (FK866 vs. control; DESeq2) for expressed protein-coding genes (baseMean ≥ 10), stratified by the compartment status of the 50-kb bin containing their TSS. B→A (n = 63) and A→B (n = 35), genes in bins called as switching by HOMER differential compartment analysis. Stable_A (n = 9,982) and stable_B (n = 1,321), genes in bins that retained their compartment assignment. Dotted line, log2FC = 0. The x-axis is restricted to the 0.5th–99.5th percentile of log2FC for display; no data were excluded (**A**). Location shift in log2FC of genes in switching bins relative to genes in stable bins of the same compartment of origin (B→A vs. stable_B; A→B vs. stable_A). Points, Hodges–Lehmann estimate of the shift; error bars, 95% confidence interval; p-values, two-sided Wilcoxon rank-sum test. Dashed line corresponds to no shift (**B**). Mean log2FC of expressed genes per 50-kb bin versus the change in compartment score of that bin (ΔE1 = mean E1 FK866 − mean E1 control). Positive E1 corresponds to the gene-dense A compartment, so positive ΔE1 indicates a shift toward A. Only bins containing at least one expressed gene are shown. Grey, stable bins; orange, switching bins. Black line, ordinary least-squares linear fit across all plotted bins with 95% confidence band; R², coefficient of determination (**C**). Compartments were derived from Hi-C first principal component (E1) values computed with HOMER at 50-kb resolution (hg19) and averaged across replicates within each condition. Bins without a confident compartment call were excluded. Genes were assigned to a single bin by their TSS (Ensembl v75, protein-coding).

**Supplemental Figure 5.**
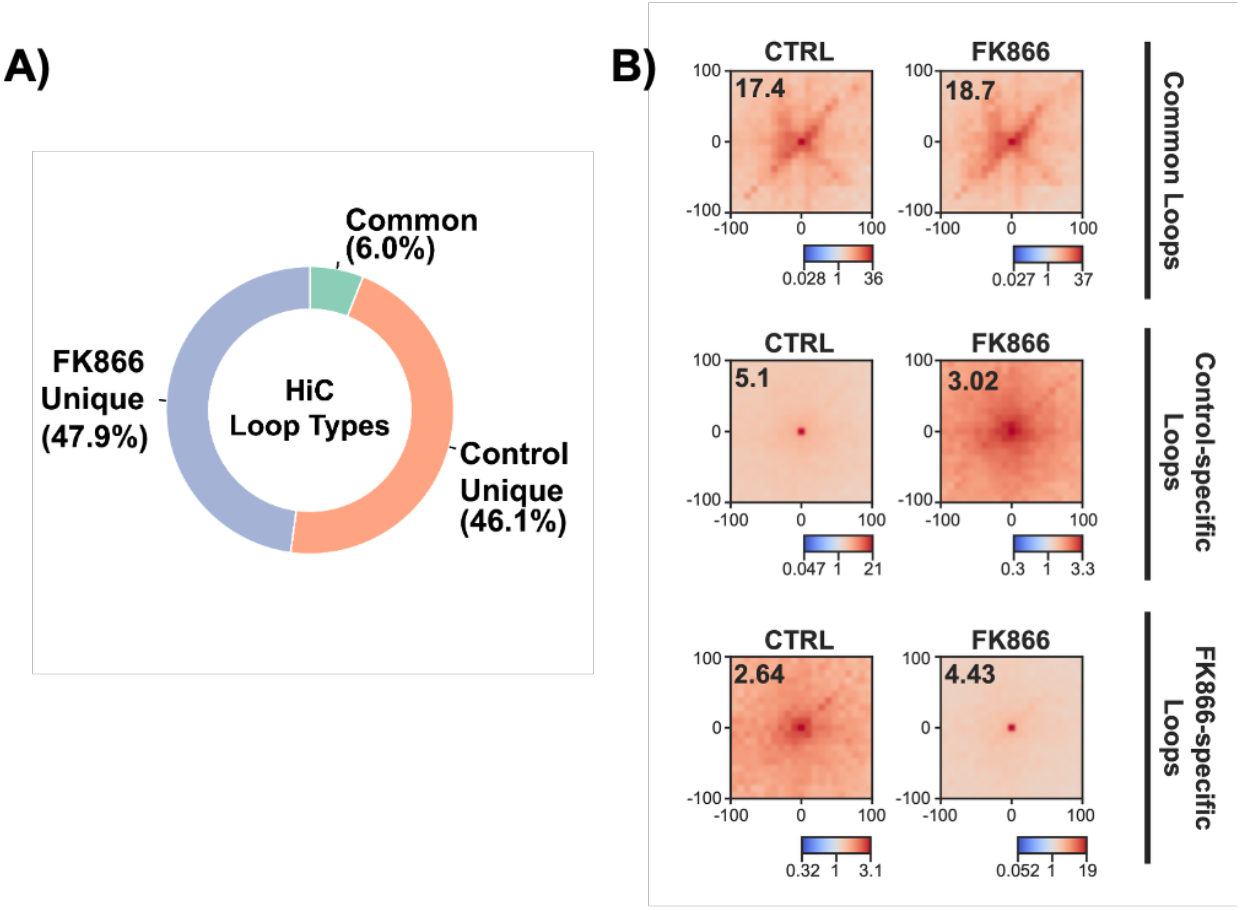
**Loops showed different frequency in Control and FK866-treated conditions by HiC assay**: Donut plot showing the percentage of loop types (common, control-only and FK866-only) detected by HiC (**A**). Aggregate Peak Analysis (APA) plots showing the enrichment of specific loops detected by HiC (**B**).

**Supplemental Figure 6.**
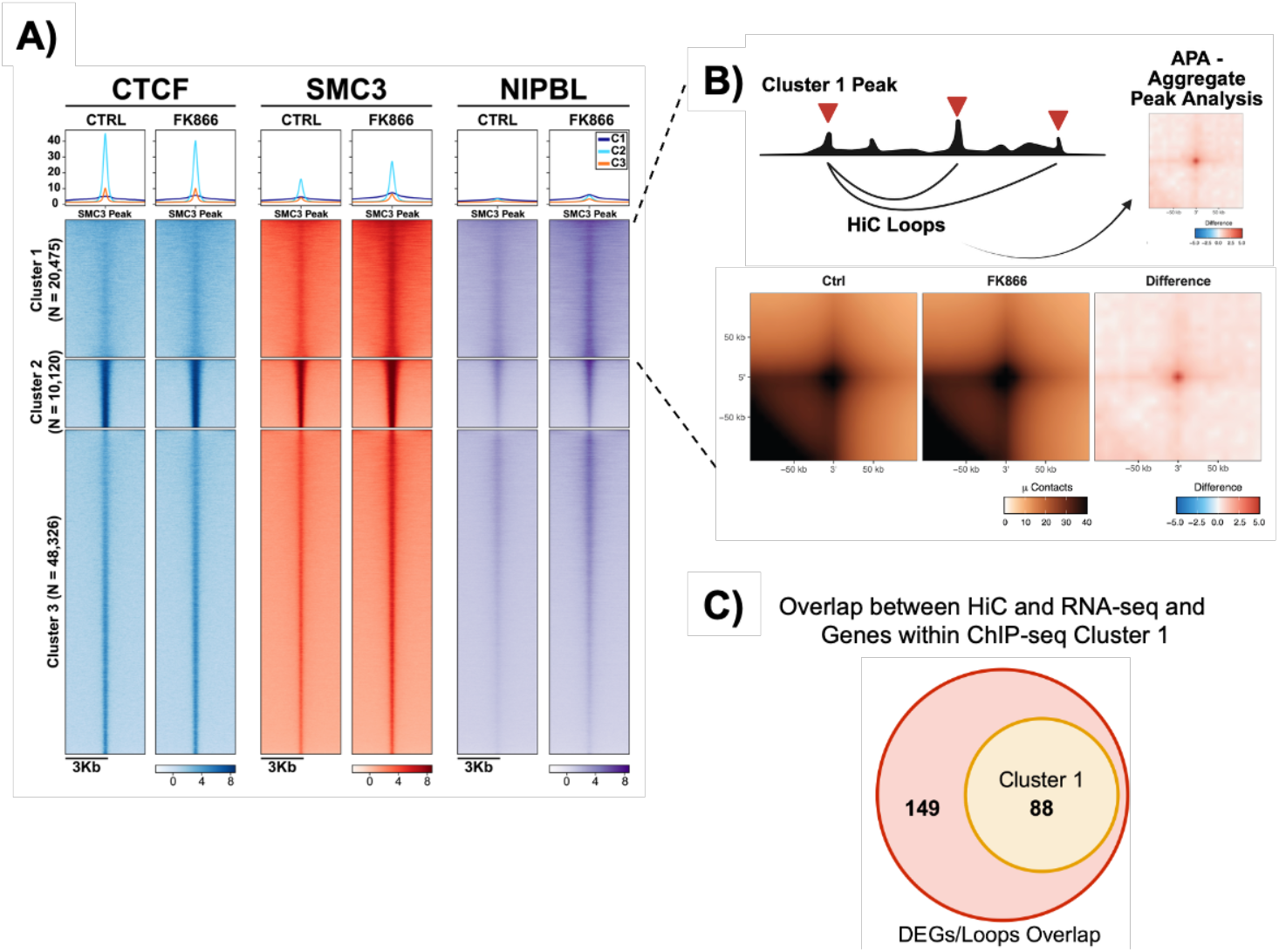
**Ectopic gain of cohesin peaks leads to increased loop extrusion**: Heatmaps showing CTCF, SMC3 and NIPBL ChIP-seq signal on SMC3 peaks. Clustering is the same as shown in Figure 4B (**A**). Aggregate Peak Analysis (APA) plots showing the enrichment of loops detected by HiC which anchors overlap peaks found in cluster 1 (**B**). Venn diagram showing the overlap of genes whose TSS was found to be overlapping with HiC loop anchors and genes annotated to ChIP-seq cluster 1 (**C**).

**Supplemental Figure 7.**
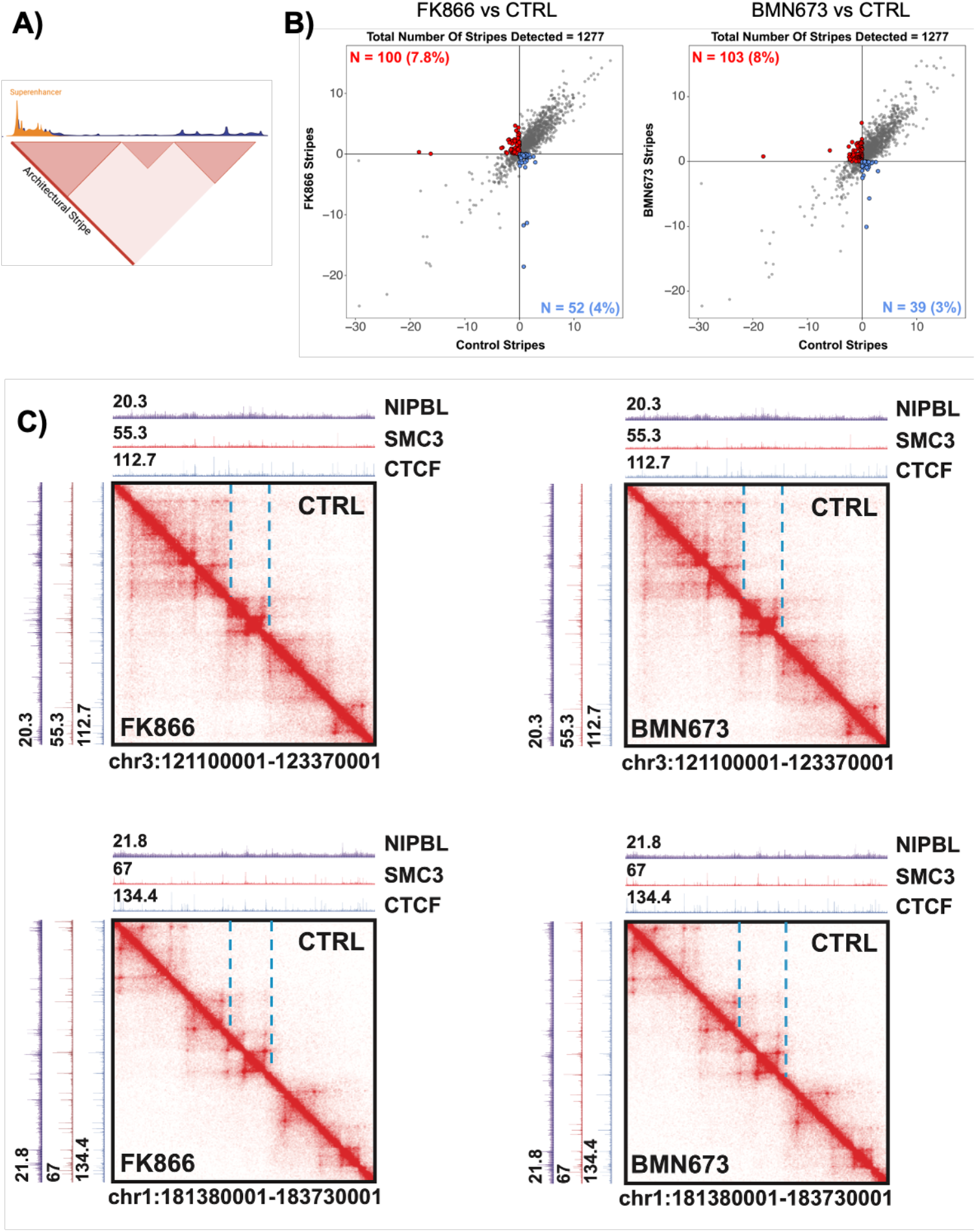
**Increased NIPBL residency time on chromatin determines formation of new stripes in FK866-treated cells**: Model of the properties of stripes (**A**). Scatterplots showing detected stripes by stripenn from TopoLink matrices (P-value < 0.05). In red and blue are indicated the lost and gained stripes upon FK866 (left) or BMN673 (right) treatment respectively (**B**). Matrices showing control (upper half) and treated (lower half) contacts. ChIP-seq read-depth normalized tracks for CTCF, SMC3 and NIPBL for the control (top) and treated (left) conditions are shown (**C**).

**Supplemental Figure 8.**
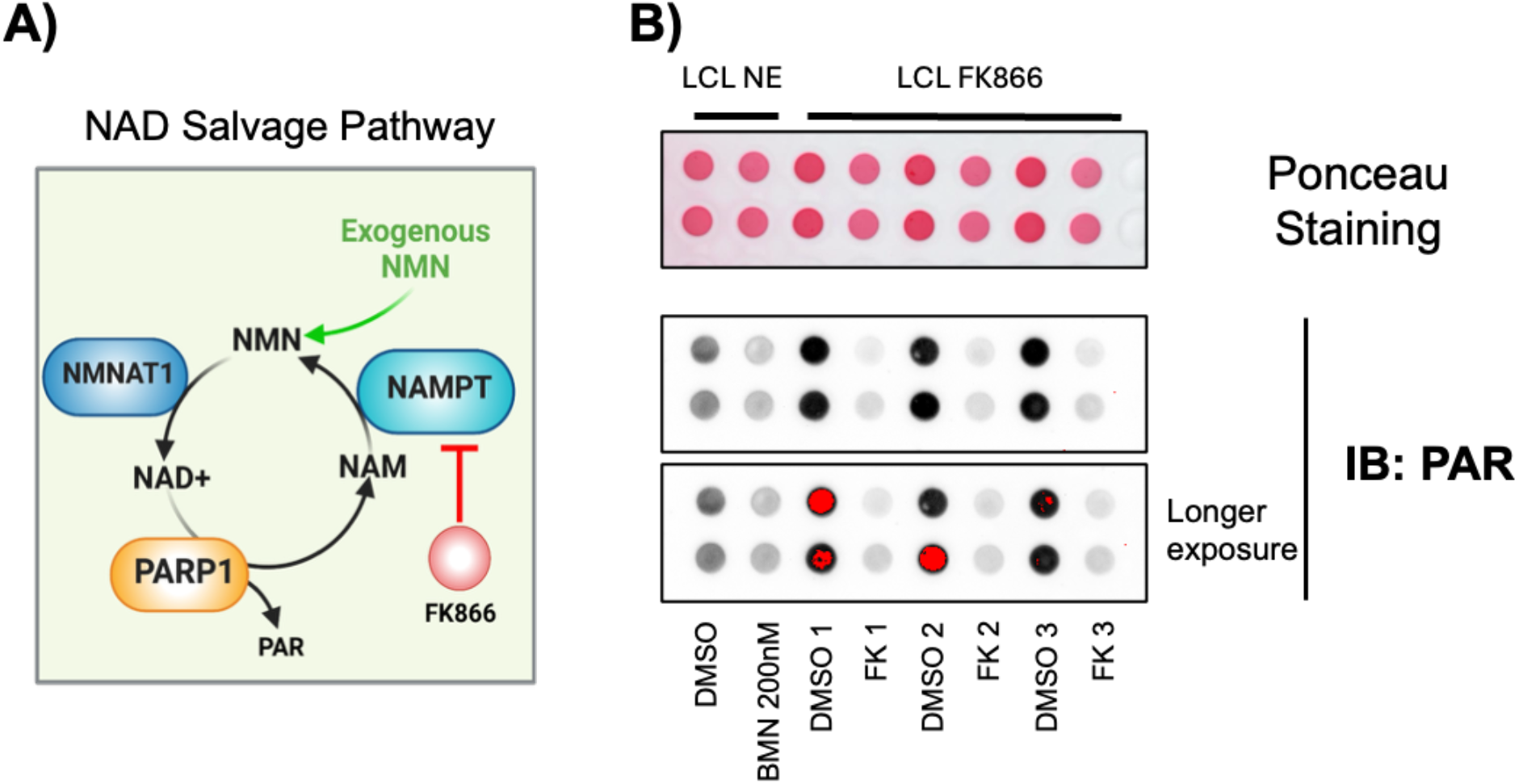
**FK866 treatment of LCLs induces a reduction in PARP1 enzymatic activity**: Model of the NAD salvage pathway (**A**). Dot Blot showing PAR levels in LCLs treated with DMSO or FK866. Protein loaded on the membrane were from either nuclear extract (NE) or whole cell lysate. Red Ponceau staining was used to quantify loaded protein level (**B**).

**Supplemental Figure 9.**
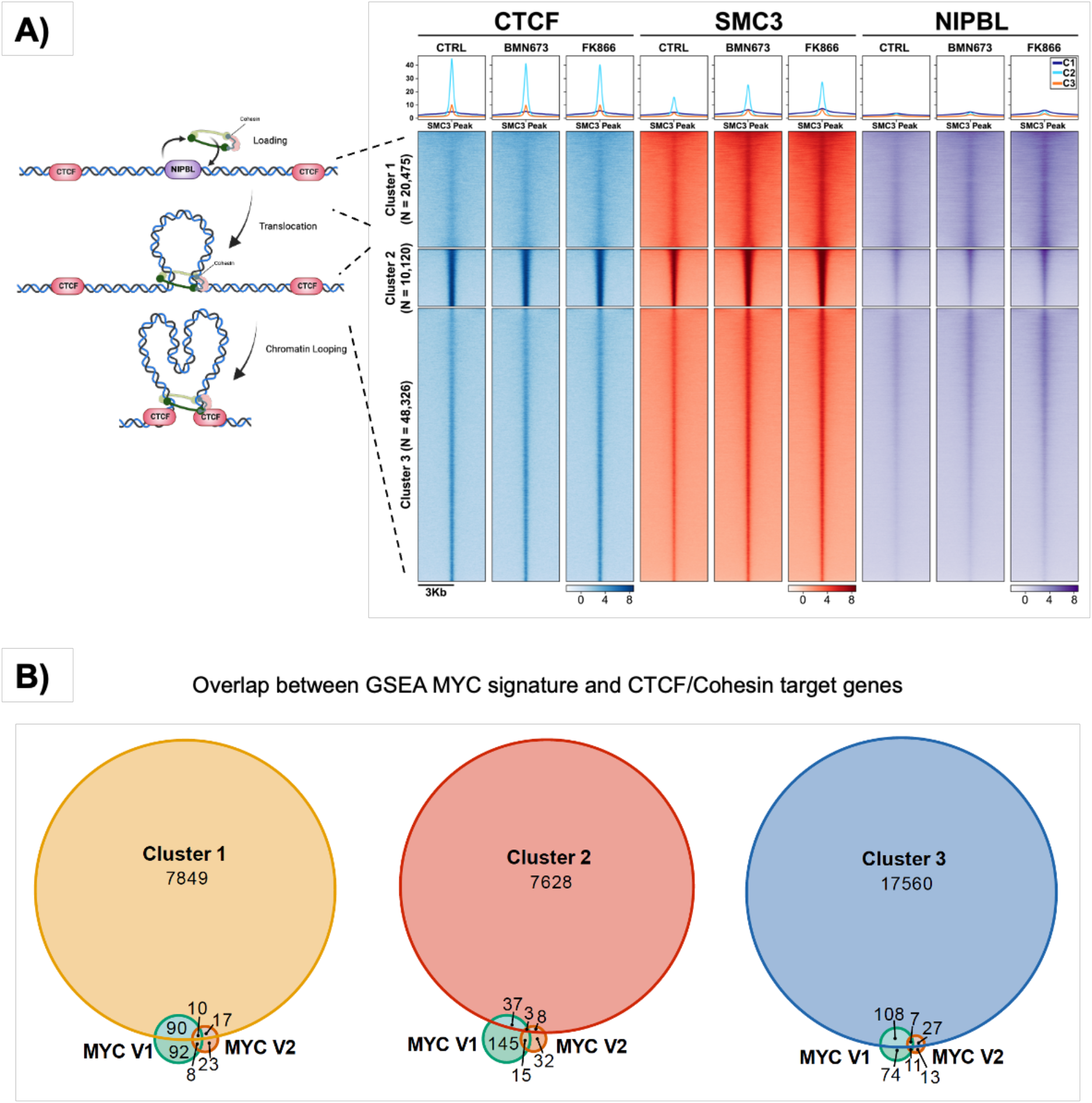
**PARP1 inhibition with BMN673 recapitulates NAMPTi**: Heatmaps showing ChIP-seq signal for CTCF, SMC3 and NIPBL on SMC3 clustered peaks in control, BMN673-treated and FK866-treated LCLs (averaged signal over N = 2 biological replicates) (**A**). Venn diagrams showing the overlaps of annotate genes in each ChIP-seq cluster and MYC GSEA pathways (V1 and V2) (**B**).

**Supplemental Figure 10.**
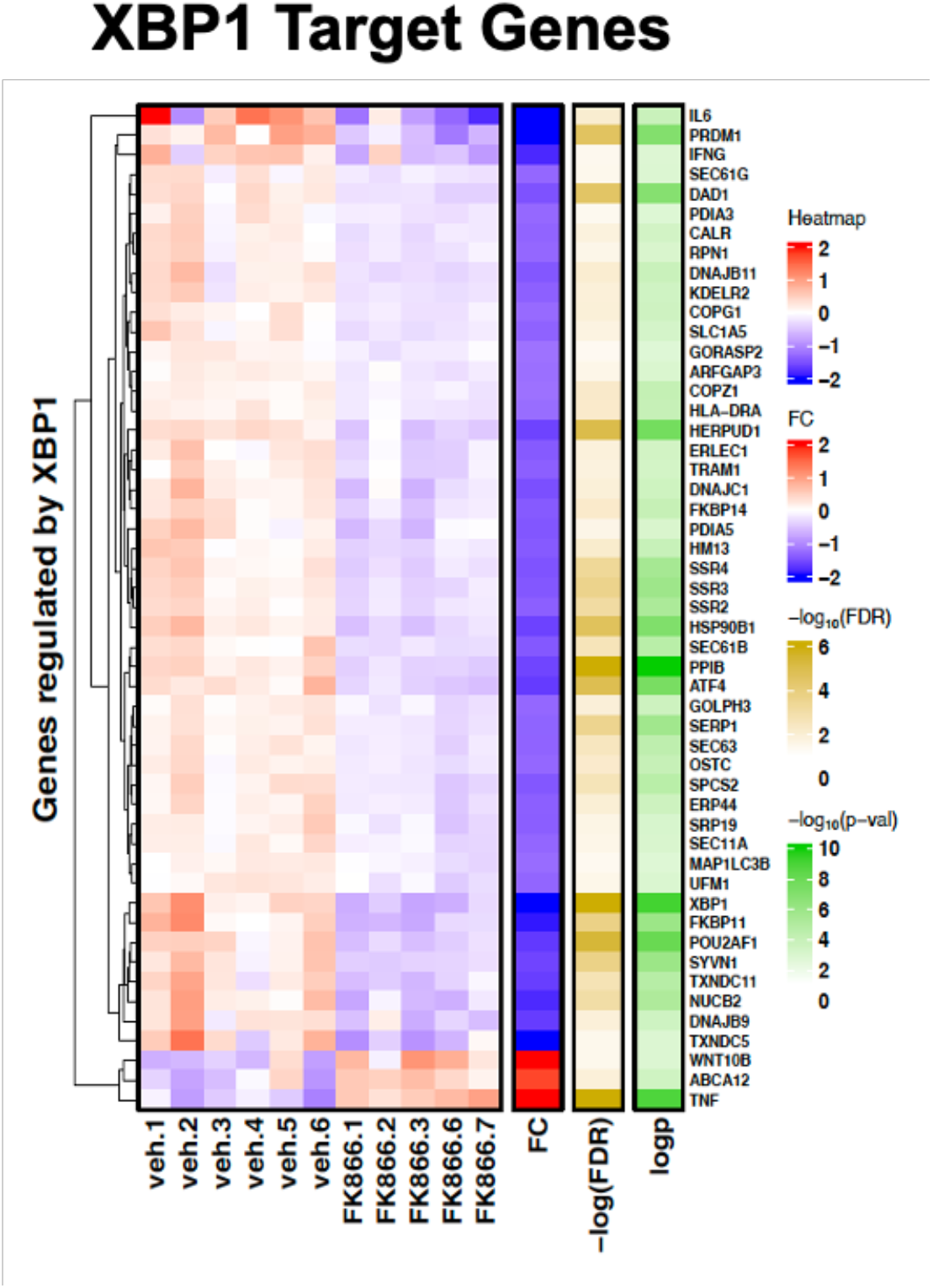
**XBP1 pathway is robustly downregulated upon NAMPTi in vivo**: Heatmap showing Z-score, log_2_FC, −log_10_(FDR) and –log_10_(P-value) of the XBP1 pathway genes.

## Notes

### Competing Interest Statement

The authors have declared no competing interest.

## References

1. Mañes-García, J., Marco-Ferreres, R. & Beccari, L. Shaping gene expression and its evolution by chromatin architecture and enhancer activity. Curr Top Dev Biol 159, 406–437 (2024).

2. Misteli, T. & Finn, E. H. Chromatin architecture is a flexible foundation for gene expression. Nat Genet 53, 426–427 (2021).

3. Wang, X. Editorial: Chromatin architecture in gene regulation and disease. Front Genet 14, 1254865 (2023).

4. Fudenberg, G., Abdennur, N., Imakaev, M., Goloborodko, A. & Mirny, L. A. Emerging Evidence of Chromosome Folding by Loop Extrusion. Cold Spring Harb Symp Quant Biol 82, 45–55 (2017).

5. Mirny, L. A., Imakaev, M. & Abdennur, N. Two major mechanisms of chromosome organization. Curr Opin Cell Biol 58, 142–152 (2019).

6. Horsfield, J. A. Full circle: a brief history of cohesin and the regulation of gene expression. The FEBS Journal 290, 1670–1687 (2023).

7. Parelho, V. et al. Cohesins functionally associate with CTCF on mammalian chromosome arms. Cell 132, 422–433 (2008).

8. Hansen, A. S. CTCF as a boundary factor for cohesin-mediated loop extrusion: evidence for a multi-step mechanism. Nucleus 11, 132–148 (2020).

9. Pentland, I. et al. Disruption of CTCF-YY1–dependent looping of the human papillomavirus genome activates differentiation-induced viral oncogene transcription. PLoS Biol 16, e2005752 (2018).

10. Ferguson, J. et al. The chromatin insulator CTCF regulates HPV18 transcript splicing and differentiation-dependent late gene expression. PLoS Pathog 17, e1010032 (2021).

11. Richman, A. et al. Multiple long-range *cis* interactions generate CTCF insulator-dependent viral chromatin domains in quiescent HSV-1 genomes. mBio e01638–25 (2025) doi:10.1128/mbio.01638-25.

12. Lang, F. et al. Author Correction: CTCF interacts with the lytic HSV-1 genome to promote viral transcription. Sci Rep 11, 5039 (2021).

13. Morgens, D. W. et al. Enhancers and genome conformation provide complex transcriptional control of a herpesviral gene. Mol Syst Biol 21, 30–58 (2024).

14. Campbell, M. et al. KSHV Topologically Associating Domains in Latent and Reactivated Viral Chromatin. J Virol 96, e00565–22 (2022).

15. Campbell, M. et al. KSHV episomes reveal dynamic chromatin loop formation with domain-specific gene regulation. Nat Commun 9, 49 (2018).

16. Kang, H., Wiedmer, A., Yuan, Y., Robertson, E. & Lieberman, P. M. Coordination of KSHV Latent and Lytic Gene Control by CTCF-Cohesin Mediated Chromosome Conformation. PLoS Pathog 7, e1002140 (2011).

17. Stedman, W. et al. Cohesins localize with CTCF at the KSHV latency control region and at cellular c-myc and H19/Igf2 insulators. EMBO J 27, 654–666 (2008).

18. Maestri, D. et al. The three-dimensional structure of the EBV genome plays a crucial role in regulating viral gene expression in EBVaGC. Nucleic Acids Research 51, 12092–12110 (2023).

19. Morgan, S. M. et al. The three-dimensional structure of Epstein-Barr virus genome varies by latency type and is regulated by PARP1 enzymatic activity. Nat Commun 13, 187 (2022).

20. Lupey-Green, L. N. et al. PARP1 Stabilizes CTCF Binding and Chromatin Structure To Maintain Epstein-Barr Virus Latency Type. J Virol 92, e00755–18 (2018).

21. Tempera, I., Klichinsky, M. & Lieberman, P. M. EBV Latency Types Adopt Alternative Chromatin Conformations. PLoS Pathog 7, e1002180 (2011).

22. Tempera, I. & Lieberman, P. M. Chromatin organization of gammaherpesvirus latent genomes. Biochimica et Biophysica Acta (BBA) - Gene Regulatory Mechanisms 1799, 236–245 (2010).

23. Guo, R. et al. MYC Controls the Epstein-Barr Virus Lytic Switch. Molecular Cell 78, 653–669.e8 (2020).

24. Patel, P. D., Alghareeb, R., Hussain, A., Maheshwari, M. V. & Khalid, N. The Association of Epstein-Barr Virus With Cancer. Cureus 10.7759/cureus.26314 (2022) doi:10.7759/cureus.26314.

25. Gewurz, B. E., Longnecker, R. & Cohen, J. I. Epstein-Barr Virus. Fields Virology. (Wolters Kluwer Health/Lippincott Williams & Wilkins, 2021).

26. Wang, C. et al. A DNA tumor virus globally reprograms host 3D genome architecture to achieve immortal growth. Nat Commun 14, 1598 (2023).

27. Lu, F. et al. EBNA2 Drives Formation of New Chromosome Binding Sites and Target Genes for B-Cell Master Regulatory Transcription Factors RBP-jκ and EBF1. PLoS Pathog 12, e1005339 (2016).

28. Arvey, A. et al. An Atlas of the Epstein-Barr Virus Transcriptome and Epigenome Reveals Host-Virus Regulatory Interactions. Cell Host & Microbe 12, 233–245 (2012).

29. Lamontagne, R. J. et al. A multi-omics approach to Epstein-Barr virus immortalization of B- cells reveals EBNA1 chromatin pioneering activities targeting nucleotide metabolism. PLoS Pathog 17, e1009208 (2021).

30. Kim, K.-D. et al. Epigenetic specifications of host chromosome docking sites for latent Epstein-Barr virus. Nat Commun 11, 877 (2020).

31. Beer, S., et al. EBNA2-EBF1 complexes promote MYC expression and metabolic processes driving S-phase progression of Epstein-Barr virus–infected B cells. Proc. Natl. Acad. Sci. U.S.A. 119, e2200512119 (2022).

32. Yang, Y. et al. Phase separation of Epstein-Barr virus EBNA2 protein reorganizes chromatin topology for epigenetic regulation. Commun Biol 4, 967 (2021).

33. Sinclair, A. J., Palmero, I., Peters, G. & Farrell, P. J. EBNA-2 and EBNA-LP cooperate to cause G0 to G1 transition during immortalization of resting human B lymphocytes by Epstein-Barr virus. EMBO J 13, 3321–3328 (1994).

34. Portal, D. et al. EBV nuclear antigen EBNALP dismisses transcription repressors NCoR and RBPJ from enhancers and EBNA2 increases NCoR-deficient RBPJ DNA binding. Proc Natl Acad Sci U S A 108, 7808–7813 (2011).

35. Maestri, D. et al. EBNA leader protein orchestrates chromatin architecture remodeling during Epstein-Barr virus-induced B cell transformation. Nucleic Acids Research 53, gkaf629 (2025).

36. Sobotka, A. A. & Tempera, I. PARP1 as an Epigenetic Modulator: Implications for the Regulation of Host-Viral Dynamics. Pathogens 13, 131 (2024).

37. Lupey-Green, L. N. et al. PARP1 restricts Epstein Barr Virus lytic reactivation by binding the BZLF1 promoter. Virology 507, 220–230 (2017).

38. Martin, K. A., Lupey, L. N. & Tempera, I. Epstein-Barr Virus Oncoprotein LMP1 Mediates Epigenetic Changes in Host Gene Expression through PARP1. J Virol 90, 8520–8530 (2016).

39. Xie, N. et al. NAD+ metabolism: pathophysiologic mechanisms and therapeutic potential. Sig Transduct Target Ther 5, 227 (2020).

40. Müller-Durovic, B. et al. A metabolic dependency of EBV can be targeted to hinder B cell transformation. Science 385, eadk4898 (2024).

41. Colevas, A. D. et al. First-in-Human Clinical Trial of a Small-Molecule EBNA1 Inhibitor, VK-2019, in Patients with Epstein-Barr-Positive Nasopharyngeal Cancer, with Pharmacokinetic and Pharmacodynamic Studies. Clin Cancer Res 31, 815–823 (2025).

42. Leibovitch, E. C. et al. Prevalence of salivary human herpesviruses in pediatric multiple sclerosis cases and controls. Mult Scler 25, 644–652 (2019).

43. Dobin, A. et al. STAR: ultrafast universal RNA-seq aligner. Bioinformatics 29, 15–21 (2013).

44. Liao, Y., Smyth, G. K. & Shi, W. featureCounts: an efficient general purpose program for assigning sequence reads to genomic features. Bioinformatics 30, 923–930 (2014).

45. Love, M. I., Huber, W. & Anders, S. Moderated estimation of fold change and dispersion for RNA-seq data with DESeq2. Genome Biol 15, 550 (2014).

46. Ramírez, F. et al. deepTools2: a next generation web server for deep-sequencing data analysis. Nucleic Acids Res 44, W160–W165 (2016).

47. Krämer, A., Green, J., Pollard, J. & Tugendreich, S. Causal analysis approaches in Ingenuity Pathway Analysis. Bioinformatics 30, 523–530 (2014).

48. 48. Raivo Kolde. pheatmap: Pretty Heatmaps. 1.0.12 10.32614/CRAN.package.pheatmap (2010).

49. Li, H. & Durbin, R. Fast and accurate long-read alignment with Burrows–Wheeler transform. Bioinformatics 26, 589–595 (2010).

50. Feng, J., Liu, T., Qin, B., Zhang, Y. & Liu, X. S. Identifying ChIP-seq enrichment using MACS. Nat Protoc 7, 1728–1740 (2012).

51. Zhang, Y. et al. Model-based Analysis of ChIP-Seq (MACS). Genome Biol 9, R137 (2008).

52. Heinz, S. et al. Simple Combinations of Lineage-Determining Transcription Factors Prime cis-Regulatory Elements Required for Macrophage and B Cell Identities. Molecular Cell 38, 576–589 (2010).

53. Ye, T. et al. seqMINER: an integrated ChIP-seq data interpretation platform. Nucleic Acids Research 39, e35–e35 (2011).

54. Wickham, H. *Ggplot2*. (Springer International Publishing, Cham, 2016). doi:10.1007/978-3-319-24277-4.

55. Grandi, F. C., Modi, H., Kampman, L. & Corces, M. R. Chromatin accessibility profiling by ATAC-seq. Nat Protoc 17, 1518–1552 (2022).

56. Danecek, P. et al. Twelve years of SAMtools and BCFtools. GigaScience 10, giab008 (2021).

57. Berest, I. et al. Quantification of Differential Transcription Factor Activity and Multiomics-Based Classification into Activators and Repressors: diffTF. Cell Reports 29, 3147–3159.e12 (2019).

58. Servant, N. et al. HiC-Pro: an optimized and flexible pipeline for Hi-C data processing. Genome Biol 16, 259 (2015).

59. Stansfield, J. C., Cresswell, K. G., Vladimirov, V. I. & Dozmorov, M. G. HiCcompare: an R-package for joint normalization and comparison of HI-C datasets. BMC Bioinformatics 19, 279 (2018).

60. van der Weide, R. H., et al. Hi-C analyses with GENOVA: a case study with cohesin variants. NAR Genomics and Bioinformatics 3, lqab040 (2021).

61. Open2C et al. Cooltools: enabling high-resolution Hi-C analysis in Python. Preprint at 10.1101/2022.10.31.514564 (2022).

62. Kramer, N. E. et al. Plotgardener: cultivating precise multi-panel figures in R. Bioinformatics 38, 2042–2045 (2022).

63. Yoon, S., Chandra, A. & Vahedi, G. Stripenn detects architectural stripes from chromatin conformation data using computer vision. Nat Commun 13, 1602 (2022).

64. Mumbach, M. R. et al. HiChIP: efficient and sensitive analysis of protein-directed genome architecture. Nat Methods 13, 919–922 (2016).

65. Lareau, C. & Aryee, M. Hichipper: A Preprocessing Pipeline for Assessing Library Quality and DNA Loops from HiChIP Data. http://biorxiv.org/lookup/doi/10.1101/192302 (2017) doi:10.1101/192302.

66. Lareau, C. A. & Aryee, M. J. diffloop: a computational framework for identifying and analyzing differential DNA loops from sequencing data. Bioinformatics 34, 672–674 (2018).

67. Ramírez, F. et al. High-resolution TADs reveal DNA sequences underlying genome organization in flies. Nat Commun 9, 189 (2018).

68. Wolff, J. et al. Galaxy HiCExplorer 3: a web server for reproducible Hi-C, capture Hi-C and single-cell Hi-C data analysis, quality control and visualization. Nucleic Acids Research 48, W177–W184 (2020).

69. Wolff, J. et al. Galaxy HiCExplorer: a web server for reproducible Hi-C data analysis, quality control and visualization. Nucleic Acids Research 46, W11–W16 (2018).

70. Lawrence, M. et al. Software for Computing and Annotating Genomic Ranges. PLoS Comput Biol 9, e1003118 (2013).

71. Sahin, M. et al. HiC-DC+ enables systematic 3D interaction calls and differential analysis for Hi-C and HiChIP. Nat Commun 12, 3366 (2021).

72. Sidoli, S., Bhanu, N. V., Karch, K. R., Wang, X. & Garcia, B. A. Complete Workflow for Analysis of Histone Post-translational Modifications Using Bottom-up Mass Spectrometry: From Histone Extraction to Data Analysis. J Vis Exp 54112 (2016) doi:10.3791/54112.

73. Yuan, Z.-F. et al. EpiProfile 2.0: A Computational Platform for Processing Epi-Proteomics Mass Spectrometry Data. J Proteome Res 17, 2533–2541 (2018).

74. Vásquez-Trincado, C. et al. Adaptation of the heart to Frataxin depletion: Evidence that integrated stress response can predominate over mTORC1 activation. Hum Mol Genet 33, 637–654 (2021).

75. Caruso, L. B. et al. PARP1 Inhibition Halts EBV+ Lymphoma Progression by Disrupting the EBNA2/MYC Axis. J Med Virol 97, e70485 (2025).

76. Hurtado-Bagès, S., Knobloch, G., Ladurner, A. G. & Buschbeck, M. The taming of PARP1 and its impact on NAD+ metabolism. Mol Metab 38, 100950 (2020).

77. Kim, M. Y., Mauro, S., Gévry, N., Lis, J. T. & Kraus, W. L. NAD+-dependent modulation of chromatin structure and transcription by nucleosome binding properties of PARP-1. Cell 119, 803–814 (2004).

78. Hulse, M., Johnson, S. M., Boyle, S., Caruso, L. B. & Tempera, I. Epstein-Barr Virus-Encoded Latent Membrane Protein 1 and B-Cell Growth Transformation Induce Lipogenesis through Fatty Acid Synthase. J Virol 95, e01857–20 (2021).

79. Wang, L. W. et al. Epstein-Barr-Virus-Induced One-Carbon Metabolism Drives B Cell Transformation. Cell Metab 30, 539–555.e11 (2019).

80. Wang, L. W. et al. Epstein-Barr-Virus-Induced One-Carbon Metabolism Drives B Cell Transformation. Cell Metab 30, 539–555.e11 (2019).

81. Wang, F. et al. Epstein-Barr virus nuclear antigen 2 specifically induces expression of the B-cell activation antigen CD23. Proc Natl Acad Sci U S A 84, 3452–3456 (1987).

82. Thorley-Lawson, D. A. & Mann, K. P. Early events in Epstein-Barr virus infection provide a model for B cell activation. J Exp Med 162, 45–59 (1985).

83. Rooney, C. M. et al. Host cell and EBNA-2 regulation of Epstein-Barr virus latent-cycle promoter activity in B lymphocytes. J Virol 66, 496–504 (1992).

84. Zhang, J. et al. LMP1 and EBNA2 constitute a minimal set of EBV genes for transformation of human B cells. Front Immunol 14, 1331730 (2023).

85. Cable, J. M., Reinoso-Vizcaino, N. M., White, R. E. & Luftig, M. A. Epstein-Barr virus protein EBNA-LP engages YY1 through leucine-rich motifs to promote naïve B cell transformation. PLoS Pathog 20, e1011950 (2024).

86. Echendu, C. W. & Ling, P. D. Regulation of Sp100A subnuclear localization and transcriptional function by EBNA-LP and interferon. J Interferon Cytokine Res 28, 667–678 (2008).

87. Ling, P. D. et al. Mediation of Epstein-Barr virus EBNA-LP transcriptional coactivation by Sp100. EMBO J 24, 3565–3575 (2005).

88. Manet, E. et al. Modulation of alternative splicing during early infection of human primary B lymphocytes with Epstein-Barr virus (EBV): a novel function for the viral EBNA-LP protein. Nucleic Acids Research 49, 10657–10676 (2021).

89. Szymula, A. et al. Correction: Epstein-Barr virus nuclear antigen EBNA-LP is essential for transforming naïve B cells, and facilitates recruitment of transcription factors to the viral genome. PLoS Pathog 15, e1007403 (2019).

90. Wang, C. et al. Viral oncogene EBNALP regulates YY1 DNA binding and alters host 3D genome organization. EMBO Rep 26, 810–835 (2025).

91. Portal, D. et al. Epstein-Barr virus nuclear antigen leader protein localizes to promoters and enhancers with cell transcription factors and EBNA2. Proc Natl Acad Sci U S A 110, 18537–18542 (2013).

92. Hong, T. et al. Epstein–Barr virus nuclear antigen 2 extensively rewires the human chromatin landscape at autoimmune risk loci. Genome Res. 31, 2185–2198 (2021).

93. Mumbach, M. R. et al. Enhancer connectome in primary human cells identifies target genes of disease-associated DNA elements. Nat Genet 49, 1602–1612 (2017).

94. Gilbert, L. A. et al. CRISPR-mediated modular RNA-guided regulation of transcription in eukaryotes. Cell 154, 442–451 (2013).

95. Hasmann, M. & Schemainda, I. FK866, a highly specific noncompetitive inhibitor of nicotinamide phosphoribosyltransferase, represents a novel mechanism for induction of tumor cell apoptosis. Cancer Res 63, 7436–7442 (2003).

96. Winkelmann, R., Sandrock, L., Kirberg, J., Jäck, H.-M. & Schuh, W. KLF2--a negative regulator of pre-B cell clonal expansion and B cell activation. PLoS One 9, e97953 (2014).

97. Winkelmann, R. et al. B cell homeostasis and plasma cell homing controlled by Krüppel-like factor 2. Proc Natl Acad Sci U S A 108, 710–715 (2011).

98. Lim, C. P., Jain, N. & Cao, X. Stress-induced immediate-early gene, egr-1, involves activation of p38/JNK1. Oncogene 16, 2915–2926 (1998).

99. Unterholzner, L. et al. IFI16 is an innate immune sensor for intracellular DNA. Nat Immunol 11, 997–1004 (2010).

100. Hamidi, T. et al. Nuclear protein 1 promotes pancreatic cancer development and protects cells from stress by inhibiting apoptosis. J Clin Invest 122, 2092–2103 (2012).

101. Jakobsen, J. S. et al. Temporal mapping of CEBPA and CEBPB binding during liver regeneration reveals dynamic occupancy and specific regulatory codes for homeostatic and cell cycle gene batteries. Genome Res 23, 592–603 (2013).

102. Schwickert, T. A. et al. Stage-specific control of early B cell development by the transcription factor Ikaros. Nat Immunol 15, 283–293 (2014).

103. Sugai, M., Gonda, H., Nambu, Y., Yokota, Y. & Shimizu, A. Role of Id proteins in B lymphocyte activation: new insights from knockout mouse studies. J Mol Med (Berl) 82, 592–599 (2004).

104. Peng, S. L., Gerth, A. J., Ranger, A. M. & Glimcher, L. H. NFATc1 and NFATc2 together control both T and B cell activation and differentiation. Immunity 14, 13–20 (2001).

105. McIntosh, M. T. et al. STAT3 imparts BRCAness by impairing homologous recombination repair in Epstein-Barr virus-transformed B lymphocytes. PLoS Pathog 16, e1008849 (2020).

106. Haarhuis, J. H. I. et al. The Cohesin Release Factor WAPL Restricts Chromatin Loop Extension. Cell 169, 693–707.e14 (2017).

107. Krantz, I. D. et al. Cornelia de Lange syndrome is caused by mutations in NIPBL, the human homolog of Drosophila melanogaster Nipped-B. Nat Genet 36, 631–635 (2004).

108. Tonkin, E. T., Wang, T.-J., Lisgo, S., Bamshad, M. J. & Strachan, T. NIPBL, encoding a homolog of fungal Scc2-type sister chromatid cohesion proteins and fly Nipped-B, is mutated in Cornelia de Lange syndrome. Nat Genet 36, 636–641 (2004).

109. Davidson, I. F. et al. DNA loop extrusion by human cohesin. Science 366, 1338–1345 (2019).

110. Kuleshov, M. V. et al. Enrichr: a comprehensive gene set enrichment analysis web server 2016 update. Nucleic Acids Res 44, W90–97 (2016).

111. Lieberman-Aiden, E. et al. Comprehensive mapping of long-range interactions reveals folding principles of the human genome. Science 326, 289–293 (2009).

112. Splinter, E. et al. CTCF mediates long-range chromatin looping and local histone modification in the beta-globin locus. Genes Dev 20, 2349–2354 (2006).

113. Barrington, C. et al. Enhancer accessibility and CTCF occupancy underlie asymmetric TAD architecture and cell type specific genome topology. Nat Commun 10, 2908 (2019).

114. Vian, L. et al. The Energetics and Physiological Impact of Cohesin Extrusion. Cell 175, 292–294 (2018).

115. Kraft, K. et al. Serial genomic inversions induce tissue-specific architectural stripes, gene misexpression and congenital malformations. Nat Cell Biol 21, 305–310 (2019).

116. Caruso, L. B. et al. PARP1 Inhibition Halts EBV+ Lymphoma Progression by Disrupting the EBNA2/MYC Axis. J Med Virol 97, e70485 (2025).

117. Shen, Y. et al. BMN 673, a novel and highly potent PARP1/2 inhibitor for the treatment of human cancers with DNA repair deficiency. Clin Cancer Res 19, 5003–5015 (2013).

118. Wainberg, Z. A. et al. Safety and antitumor activity of the PARP inhibitor BMN673 in a phase 1 trial recruiting metastatic small-cell lung cancer (SCLC) and germline BRCA-mutation carrier cancer patients. JCO 32, 7522–7522 (2014).

119. Bjornevik, K. et al. Longitudinal analysis reveals high prevalence of Epstein-Barr virus associated with multiple sclerosis. Science 375, 296–301 (2022).

120. 120. Singh, D. R., et al. Epstein–Barr virus LMP1 protein promotes proliferation and inhibits differentiation of epithelial cells via activation of YAP and TAZ. Proc. Natl. Acad. Sci. U.S.A. 120, e2219755120 (2023).

121. Sun, Y. et al. Regulation mechanism of EBV-encoded EBER1 and LMP2A on YAP1 and the impact of YAP1 on the EBV infection status in EBV-associated gastric carcinoma. Virus Res 343, 199352 (2024).

122. Van Sciver, N. et al. Hippo signaling effectors YAP and TAZ induce Epstein-Barr Virus (EBV) lytic reactivation through TEADs in epithelial cells. PLoS Pathog 17, e1009783 (2021).

123. Wen, F. et al. Epstein-Barr virus infection upregulates extracellular OLFM4 to activate YAP signaling during gastric cancer progression. Nat Commun 15, 10543 (2024).

124. Maestre, L. et al. Expression pattern of XBP1(S) in human B-cell lymphomas. Haematologica 94, 419–422 (2009).

125. Zhu, H. et al. Ufbp1 promotes plasma cell development and ER expansion by modulating distinct branches of UPR. Nat Commun 10, 1084 (2019).

126. Tang, T. F. et al. Regulatory network of BLIMP1, IRF4, and XBP1 triad in plasmacytic differentiation and multiple myeloma pathogenesis. Cellular Immunology 380, 104594 (2022).

127. Todd, D. J. et al. XBP1 governs late events in plasma cell differentiation and is not required for antigen-specific memory B cell development. J Exp Med 206, 2151–2159 (2009).

